# Non-cell-autonomous HSC70.1 chaperone displays homeostatic feed-back regulation by binding its own mRNA

**DOI:** 10.1101/2022.05.10.491294

**Authors:** Lei Yang, Yuan Zhou, Shuangfeng Wang, Ying Xu, Steffen Ostendorp, Melissa Tomkins, Julia Kehr, Richard J. Morris, Friedrich Kragler

**Affiliations:** Max-Planck-Institute of Molecular Plant Physiology, Wissenschaftspark Golm, Am Mühlenberg 1, 14476 Golm, Germany; Universität Hamburg, Institute for Plant Science and Microbiology, Ohnhorststr. 18, 22609 Hamburg, Germany; Computational and Systems Biology, John Innes Centre, Norwich, NR4 7UH UK

## Abstract

Heat shock proteins of the HSC70/HSP70 family are evolutionarily conserved chaperones that are involved in protein folding, protein transport and RNA binding. Arabidopsis HSC70 chaperones are thought to act as housekeeping chaperones and as such are involved in many growth-related pathways. Whether Arabidopsis HSC70 binds RNA and its function has remained an open question. Here, we show that the HSC70.1 chaperone binds its own mRNA via its C-terminal Short Variable Region (SVR) and inhibits its own translation. We propose that this negative protein-transcript feedback loop may establish an on-demand chaperone pool that allows for a rapid response to stress. Furthermore, we show that the SVR encoding RNA region is necessary for *HSC70*.*1* transcript mobility to distant tissues and that *HSC70*.*1* transcript and not protein mobility is required to rescue root growth and flowering time of *hsc70* mutants. In summary, it seems that the Arabidopsis HSC70.1 chaperone can form a complex with its own transcript to regulate its translation and that both protein and transcript can act in a non-cell-autonomous manner maintaining chaperone homestasis between tissues.

## INTRODUCTION

Heat shock proteins of the HSC70/HSP70 family are evolutionarily conserved chaperones that assist folding and formation of functional structural domains of client proteins. By this means HSC70 chaperones contribute to environmental and genomic stress tolerance (Mayer & Bukau, 2005; Noël *et al*., 2007). In addition, HSC70s are often found to facilitate protein import into cellular compartments. Similarly, some plant viruses produce HSC70-related chaperones that interact with viral RNA-protein (RNP) complexes and intercellular channels to enable viral transport to neighboring cells (Gilbertson & Lucas, 1996; Lazarowitz & Beachy, 1999; Medina *et al*., 1999; Alzhanova *et al*., 2001), suggesting a potential role for HSC70-related chaperones in intercellular transport of macromolecules. Yeast and human HSC70 chaperones are reported to bind to 3’ UTRs AU-rich motifs present in distinct mRNAs either via their conserved ATPase-and substrate – binding domains or their C-terminal domain (Zimmer *et al*., 2001; Kishor *et al*., 2013; Kishor *et al*., 2017). Although strong evidence exists that HSC70 family members can bind to mRNAs the function of this RNA-interaction beside that it might stabilize interacting transcripts remains elusive as RNA bound HSC70 chaperones show no changed *in vitro* protein folding activity (Malter, 1989; Chen & Shyu, 1995; Henics *et al*., 1999; Guhaniyogi & Brewer, 2001; Kishor *et al*., 2017).

Interestingly, HSC70 chaperones are also abundant in phloem exudates (Aoki *et al*., 2002; Giavalisco *et al*., 2006) and in plasmodesmal cell fractions (Aoki *et al*., 2002). Based on these observation, we hypothesised that the HSC70 chaperone may play a role in macromolecular movement between cells and via the phloem vasculature. Further studies led to the proposal that graft-mobile mRNAs may form phloem-specific RNP complexes consisting of PHLOEM PROTEIN 16 (PP16), RNA BINDING PROTEIN 50 (RBB50; related to pyrimidine tract binding proteins), elF-5A elongation factor, TRANSLATIONALLY CONTROLLED TUMOR PROTEIN1 (TCTP1), and HSC70 (Ham *et al*., 2009; Saplaoura & Kragler, 2016). The PP16 / RBP50/ TCTP1 / HSC70 complex was suggested to bind to polypyrimidine track binding (PTB) RNA motifs (UUCUCUCUCUU) found in many 3’ UTRs and to facilitate long-distance RNA transport via the phloem (Ham *et al*., 2009; Yang *et al*., 2019). In line, the individual PP16, RBP50, and HSC70 phloem proteins were reported to move between cells in single-cell microinjection assays (Xoconostle-Cázares *et al*., 1999; Aoki *et al*., 2002; Ham *et al*., 2009). A more detailed analysis of pumpkin HSC70s revealed a structural motif in the less conserved HSC70 C-terminal Short Variable Region (SVR) domain, enabling HSC70 chaperones to move to neighboring pumpkin cells after microinjection into single cells (Aoki *et al*., 2002). However, a functional role of this complex in phloem-mediated RNA transport and direct interaction of HSC70 with mobile transcripts remains to be established.

In higher plants, several hundred mRNAs were reported to move over heterologous graft junctions to distant body parts. Common mobile transcripts were identified in distantly related plant species such as arabidopsis, cucumber, watermelon, grapevine, *Nicotiana benthamiana*, soybean, and tomato (Notaguchi *et al*., 2015; Thieme *et al*., 2015; Yang *et al*., 2015; Walther & Kragler, 2016; Zhang, Z *et al*., 2016; Xia *et al*., 2018; Li *et al*., 2021), among this set of conserved graft-mobile mRNAs are *HSC70* transcripts. *A. thaliana HSC70*.*1* transcript was shown to move shoot-to-root in juvenile plants. Notably, *HSC70*.*1* transport depends on secondary ribonucleic m^5^C modifications (Yang *et al*., 2019), suggesting that *HSC70*.*1* transcript transport to distant tissues may depend on distinct RNA structures, as seen with other mobile protein-encoding transcripts (Zhang, W *et al*., 2016; Kehr & Kragler, 2018; Yang *et al*., 2019).

To characterise a potential chaperone-RNA interaction, we identified HSC70.1 interacting mRNAs. We show that the HSC70.1 C-terminal SVR sequence acts as an RNA binding domain that can interact with its own transcript. Both *in vivo* and *in vitro* translation assays suggest that HSC70.1 binding to its encoding mRNA prevents its own translation, thus establishing a negative translational feedback loop. To address the potential redundant function of graft mobile *HSC70* mRNA and protein in receiving tissues, we asked whether *A. thaliana* HSC70.1 transcript and protein transport depends on distinct RNA or protein motif(s) and whether transcript or protein mobility is required for normal plant growth. Our results indicate that *A. thaliana HSC70*.*1* transcript movement over graft junctions depends on the same region encoding the C-terminal SVR RNA binding motif. In contrast to HSC70.1 protein, *HSC70*.*1* transcript mobility is required for complementing flowering and root growth phenotypes.

## MATERIAL AND METHODS

### Plant Material and Growth

*Arabidopsis thaliana* (Columbia-0) seeds and respective transgenics and mutants were used in the grafting and complementation experiments. Seeds were sterilized with 70% ethanol, 1% Tween 20 for 5 minutes, and 4% sodium hypochlorite for 5 minutes, washed 5 times with sterile water and resuspend in 0.15% selected agar prior transfer to plates. For root growth measurements seeds were placed and germinated on vertical half-strength MS medium plates (0.5 MS salts, 1% sucrose, and 1% micro agar [Duchefa Biochemie]) in controlled growth chambers (Percival) under long day light conditions (16 h light / 8 h dark; day 22°C / night 19°C; light intensity: 170 μE·m^−2^·s^−1^). For rosette leaves growth and flowering time measurement, seeds were germinated directly on soil and transferred 10 days after germination (DAG) to single pots and grown under long day light conditions (16 h light / 8 h dark; day 22°C / night 19°C; light intensity: 170 μE·m^−2^·s^−1^; relative humidity: 60%). The *hsc70-1* (SALK_135531) and *hsp70-4* (SALK_088253) T-DNA insertion mutants were obtained from the Arabidopsis Biological Resource Center at Ohio State University (Alonso *et al*., 2003). *hsc70-1 hsp70-4* mutant seeds was obtained from Su’s lab in Fudan University, China (Leng *et al*., 2017).

### *Arabidopsis thaliana* Hypocotyl Grafting

*Arabidopsis thaliana* seeds were vertically grown on plates (0.5 MS salts, 1% sucrose, and 1% micro agar [Duchefa Biochemie]) in short day light conditions (8 h light / 16 h dark; day 22°C / night 19°C; light intensity: 170 μE·m^−2^·s^−1^). Seedlings (7-8 DAG) with ∼ 4 cm long hypocotyls were cut in the upper half of the hypocotyl using a sterile razor blade and silicon micro-tubes with 0.3 mm internal diameter were used to stabilize the graft junction. Grafted plants were transferred to new plates and vertically grown (0.5 MS salts, 1% sucrose, and 1% micro agar [Duchefa Biochemie]) in short day light conditions. After grafting (6 - 7 days), adventitious roots appearing on the upper hypocotyl junction were removed every day. 10 days after grafting successful grafted plants were analyzed by confocal laser scanning microscopy. 14 days after grafting plants were transferred to liquid culture (0.5 MS salts, 1% sucrose), prior RNA isolation at 30 days after grafting or to soil for flower time measurements.

### Expression Constructs

*35S::YFP-HSC70* fusion constructs were produced by PCR amplifying *HSC70*.*1* full-length CDS from cDNA templates (FK1527/FK1528) and cloning it into the pENTR4 vector (Addgene) using Bam HI and EagI sites. The *HSC70*.*1* CDS fragment confirmed by sequencing was transferred into pEarlygate104 binary vector (Earley *et al*., 2006) by LR recombinase GW reaction. The *HSC70*.*1 ΔSVR* (FK1527/FK1529) and *SVR* (FK1530/FK1528) constructs were produced using the same strategy. *HSC70*.*1M* and *HSC70*.*1M ΔSVR* codon usage modified constructs were created using a synthesized DNA PCR template to replace the 3’ end of wild type *HSC70*.*1* 3’ CDS to produce *HSC70*.*1M* (modified CDS base 1185 to 1956) and *HSC70*.*1M ΔSVR* (modified CDS base 1185 to 1812) (for sequences see Supplementary Dataset 1) by using existing *BgIII* and *EagI* restriction sites into pENTR4 HSC70.1 vector. Both *HSC70*.*1M* CDS constructs were transferred into pEarlygate104 binary vector by LR recombinase reaction. The resulting binary vectors were introduced into the Agrobacterium strain *AGL1* and transformed into *A. thaliana* Col-0 and *hsc70*.*1 hsp70*.*4* plants using the double floral dip method (Davis *et al*., 2009).

### RNA Isolation

Grafted scion and root tissues were separately harvested into 2 ml tubes containing metal beats and frozen in liquid nitrogen. After grinding the broken tissue was supplemented with 0.75 mL Trizol LS Reagent (Invitrogen), homogenized (20 seconds vortexing), and incubated for 5 minutes at RT to permit complete dissociation of nucleoprotein complexes. After adding 0.2 mL of chloroform per 0.75 mL of TRIzol™ LS Reagent used for lysis, samples were vortexed vigorously and incubated for 5 minutes at RT, and centrifuged for 15 minutes (12,000 × g at 4°C). The appearing upper aqueous phase was transferred to 1.5 ml RNase-free tubes and supplemented with 1 volume isopropanol and 1/10 volume 3M NaOAc (pH 5.2), inverted 3 times, and incubated at -20°C overnight. After centrifugation for 30 minutes at 4°C and the RNA pellets were washed two times with 1 mL ice-cold 80% ethanol, one time with 99% ethanol, dissolved in 10-25 μL DEPC H_2_O, incubated 5 min at 65°C, and stored at -80°C till further use.

### Reverse Transcription Reactions and RT-PCR

Reverse transcription reaction was performed using a reverse transcription kit (Promega). First, total RNA (∼1.5 μg) and an oligo (dT) mixture were denatured at 70°C for 10 minutes and annealed for 5 minutes at 37°C, submitted to RT-reaction (42°C, 90 minutes), and to deactivation (72°C, 10 minutes). RT-PCR was conducted under standard PCR conditions after testing cDNA samples quality using *ACTIN2* (FK424/FK425) primers (28 cycles). For detection of mobile mRNA by RT-PCR, 45 PCR cycles (for presence) or 50 PCR cycles (for absence) using specific PCR primers were applied (Supplementary Dataset 2).

### Quantitative RT-PCR assay

To measure transcript levels by qRT-PCR we used the SYBR Green method and an ABI System Sequence Detector (Applied Biosystems 7900HT fast Real-time PCR). For all assays three technical replicates were performed. The regiment of thermal cycling was as follows: Step 1: 1 cycle, 2 minutes at 50°C; Step 2: 1 cycle, 10 minutes at 95°C; Step 3: 40 cycles, 15 seconds at 95°C, 1 minutes at 60°C. Dissociation step: 15 s at 95°C, 15 s at 60°C, 15 s at 95°C. For specific PCR primers used see Dataset EV2.

### Microscopy

To detect YFP-HSC70.1 fusions in hypocotyl-grafted *A. thaliana* plants (10 days after grafting) we used a Leica SP5 or SP8 Confocal Laser Scanning Microscope (CLSM). To detect low abundant YFP fluorescent signals we used a CLSM equipped with HyD hybrid detector (Leica SP8). Z-stack images were assembled and processed using the Fiji software package as described (Yang *et al*., 2019).

### Protein Isolation and Immunoblot Detection

Ten-day-old plants (for each replicate approx. 300 - 360 plants) expressing YFP-HSC70.1, YFP- HSC70.1 ΔSVR, YFP-HSC70.1 SVR, YFP-HSC70.1M, and YFP-HSC70.1M ΔSVR were harvested for protein isolation. Frozen and ground samples were transferred to 1.5. mL tubes, incubated with 0.2 mL protein extraction buffer (50 mM Tris-HCl pH 8.0, 2.5 mM EDTA, 150 mM NaCl, 10% glycerol, 0.5 mM PMSF, 1 Tablet/10 mL EDTA free Protease inhibitor cocktail, Roche), vortexed vigorously, incubated on ice for 10 minutes, centrifuged 10 minutes (12,000 × g at 4°C), transferred to a new 1.5 mL RNase-free tube, and SDS loading buffer was added prior incubating the samples at 95°C (5 minutes) and samples were submitted SDS-PAGE. To detect YFP fusion proteins the samples were separated on 10% SDS-PAGE gels and transferred to nitrocellulose membranes. After blocking (5% w/v milk TBS-T solution) and incubation with 1:10,000 anti-GFP from rat (Roche) primary antibody, washing, and incubation with 1:20,000 anti-rat IgG Horseradish Peroxidase (HRP) secondary antibodies (Promega) the chemiluminescent (ECL™ Prime Western Blotting Detection Reagent, GE Healthcare) signal was detected using the ChemiDoc MP Imaging system (Bio-Rad). Coomassie blue S-stained RuBisCO large subunit served as a loading control.

### RNA-Immunoprecipitation (RIP)

*YFP-HSC70*.*1, YFP-HSC70*.*1 ΔSVR, YFP-HSC70*.*1 SVR* and *YFP-HSC70*.*1M* transgenic seeds were vertically grown on plates under short day conditions for 10 days and then subjected to irradiation with 254 nm UV light at a dose of 500 mJ /cm2 (UVP CL-1000 UV crosslinker) on ice and frozen in liquid nitrogen immediately after crosslinking. Ground samples were supplemented with 1 mL lysis buffer (150 mM NaCl, 50 mM Tris-HCL pH7.5, 5 mM MgCl_2_, 10% glycerol, 1% NP-40 (IGEPAL), 0.5 mM DTT, 1 mM PMSF, and 1% plant PIC (Gold Bio), incubated on ice for 30 minutes and gently mixed for lysis. After centrifugation (15 minutes at 12,000 × g at 4°C) 1 mL of the supernatant was transferred to new tubes with 10 μL RNase inhibitor (Promega) and 25 μL beads (equilibrated GFP-Trap® M beads; Chromotek) and incubated (gentle end-over-end mixing) at 4°C for 2 h. The supernatants were removed from magnetically separated beads (inverted six times) and washed three times with 1 mL ice-cold wash buffer (150 mM NaCl, 50 mM Tris-HCL pH7.5, 5 mM MgCl_2_, 0.5 mM DTT). RNA was isolated using the Trizol protocol (see above) and submitted to RT-PCR (see above).

### *In vitro* Transcription

YFP and YFP-HSC70.1 DNA fragments for *in vitro* production of RNA were produced by PCR using T7 YFP FP/ T7 YFP RP and T7 YFP-HSC70.1 FP/ T7 YFP-HSC70.1 RP primers using pEarlyGate104 HSC70.1 plasmid as template. *YFP-HSC70*.*1* and *YFP* RNA were produced (100 μg) using T7 RNA transcription kit (Promega P1320) according to the manufacturer’s instructions. The RNAs were analyzed by 1% agarose gel electrophoresis to calculate the concentration and confirm the absence of degradation. For Microscale Thermophoresis, HSC70.1 and BAG1 protein encoding sequences were amplified from cDNAs with an additional T7 promotor sequence at the 5’ ends. 10 pmol of DNA were used as template per 100 μl RNA synthesis reaction. *In vitro* RNA synthesis was performed as described for 3 h at 37°C (Cazenave and Uhlenbeck, 1994) and subsequently purified using the RNA clean & concentrator-25 kit (Zymo Research).

### RNA-binding Quantification by Microscale Thermophoresis

RNA binding quantification of YFP-HSC70.1 and YFP protein was performed with cell lysates from 14 day-old transgenic *Arabidopsis thaliana* plants. Cell extracts from YFP-HSC70.1 and YFP expressing lines were made freshly as described previously with minor modifications (Chen et al., 2017). In brief, plant material was ground in liquid nitrogen using mortar and pestle. 200 μL of 2-fold MST buffer (100 mM HEPES pH 7.5, 300 mM NaCl, 20 mM MgCl2, 0.2 % (v/v) NP-40, 2 mM PMSF, 2 mM AEBSF, 0.5 U/μL RiboLock) were added per 100 mg of ground plant material and incubated on ice for 5 minutes. Cell lysates were centrifuged twice at 13,000 g for 10 minutes at 4°C and diluted 1:1 with water. Dilutions were made to achieve fluorescence reads between 200 and 1600 counts. For RNA binding quantification, serial dilutions of target RNAs were made and assays were performed according to the manufacturer’s instructions. Samples were measured in standard capillaries on a Monolith NT.115 (NanoTemper GmbH) and analyzed using MO.Analysis software and GraphPad Prism 5. Binding was regarded as true when a signal-to-noise ratio and response amplitude larger than 5 was achieved.

### Translation Assays

YFP-HSC70.1 protein was enriched from protein extract obtained from transgenic *35S::YFP-HSC70*.*1* seedlings using anti GFP/YFP AB beads following the RNA-Immunoprecipitation experiment (see description above) without crosslinking treatment. YFP-HSC70.1 beads were resuspended in 100μl BSA (0.5 μg/μL) solution. *In vitro* translation assays were performed at 25°C for 120 minutes using Wheat Germ Expression (WGE) kit (Promega: L4380) and detection of produced protein was performed using the FluoroTect™ Green_Lys_ tRNA (Promega: L5001) labeling system. Note that no, or barely detectable YFP-HSC70.1 translation was observed using the standard WGE reaction protocol provided by the manufacturer. After optimization of *YFP-HSC70*.*1* RNA template, potassium acetate, and magnesium acetate concentrations, we were able to detect YFP-HSC70.1 protein translation. The optimized *in vitro* translation reaction with a final volume of 50 μL contained 25 μL wheat germ extract, 4 μL complete amino acid mixture, 2 μg *YFP-HSC70*.*1* (4 μl) or *YFP* RNA (1.5 μl), 5 μL potassium acetate (1 M), 1 μL magnesium acetate (50 mM), 1 μL RNasin ribonuclease inhibitor (40 U/μL), 2 μL FluoroTect™ Green_Lys_ tRNA, and increasing YFP-HSC70.1 protein (0.07, 0.14, 0.28, 0.56 or 1.12 μg bound on AB beads resuspended in 0.5 μg/μL BSA). The translation reactions were terminated by placing on ice and 10 μL samples were used for SDS-PAGE and western blot detection / fluorescence detection using a Typhoon FLA 7000 imaging system with a 532 nm excitation following the FluoroTect^™^Green_Lys_ *in vitro* Translation Labeling System instructions provided by the supplier (Promega, L5001).

### *In Vivo* HSP70 Inhibition

YFP-HSC70.1, YFP-HSC70 ΔSVR, and YFP transgenic plants (14 days after germination on 0.5 MS plates supplemented with 1% sucrose; n>20) were transferred to 5 mL liquid 0.5 MS medium with 1% sucrose and incubated for 15, 30, and 60 minutes with 150 μM HSP70 inhibitor VER-155008 (Sigma SML0271; resuspended in DMSO) or 0.1% DMSO (mock treatment) at 20°C. After 15, 30, and 60 minutes protein samples (150 mg) from three independent replicates for each construct and time point were collected for western blot (1:10,000 anti-GFP from rat, Roche; see methods above) and RNA samples (150 mg) for qRT PCR (YFP specific primers; see methods above). The total protein loaded for western blot assays was adjusted according to coomassie staining and YFP fusion protein detection was performed by chemiluminescent (ECL™ Prime Western Blotting Detection Reagent, GE Healthcare; see methods above) and was quantified by chemiluminescent emission detected by a Bio-Rad ChemiDoc MP Imaging system. The Image Lab software (Version 5.2.1) Quantity Tools function was used to calculate the relative (non-saturated) density of protein bands.

### Biolistic bombardment

Leaves of 36 days old *N. benthamiana* and *N. sylvestris* plants were used for biolistic bombardment of DNA-coated 1 μm Gold Microcarriers (Bio-Rad) with a Biolistic® PDS-1000/He Particle Delivery System (Bio-Rad) as described (Winter *et al*., 2007). To prepare gold microcarriers 60 mg of gold particles were washed three times with 70% ultra-pure ethanol and resuspend in 1 mL ultra-pure ddH20. 25 μl of gold microcarriers (60 mg/ml) were mixed with binary plasmid DNA (∼7 nmol) (*YFP-HSC70*.*1, YFP-HSC70*.*1 ΔSVR, YFP-HSC70*.*1 SVR*, or *YFP)*, 25 μL 2.5 M CaCl_2_, vortexed for 2 minutes, mixed with 20 μL 0.1 M spermidine, submitted to washing steps, and used for biolistic bombardment using the following parameters: Helium pressure 1100 psi (7584.2 kPa); Vacuum of 27-28.0 Hg and examined under a CSLM SP8 (Leica) 36-48h after bombardment for presence of YFP fluorescence in neighboring cells as described (Winter *et al*., 2007).

### RNAseq analysis of RIP

Total RNA from three replicated RIP samples from erYFP and HSC70.1-GFP transgenics was submitted to Ilumina cDNA library production and sequenced (± 50 million reads paired-end RNAseq; BGI, China). The RNAseq data sets were analyzed against all annotated *A. thaliana* CDS (Araport 11 release) using the CLC Genomics Workbench v21 software (Quiagen) using default settings except that a similiarity faction of 0.9 was set. Enrichment of log2 > 1 and FDR p value ≦ 0.05 in the bulk HSC70.1 RIP sample vs. the bulk erYFP sample RNAseq reads were considered as significant as presented in Table S1.

### Computational simulation of HSC70 feedback loop

We first set up equations to describe the *in vitro* translation experiments (Figs 2C and D). To represent the experimental conditions, we assumed a constant amount of HSC70.1 mRNA, ribosomal proteins and associated machinery. Protein degradation was not included. A baseline translation rate for mRNA in the absence of HSC70.1 protein was set. To mimic the enhancement of ribosomal activity in the presence of chaperones, we allowed for the translation rate to increase with HSC70.1 protein concentration following a Hill equation. Binding of HSC70.1 mRNA to its own protein (Fig 1D) was assumed to block its translation. Different amounts of HSC70.1 protein were added to the system and the amount of translated HSC70.1 mRNA was computed by solving the set of ordinary differential equations derived from the described interactions. After ensuring the model was able to qualitatively reproduce the experimental observations shown in Fig 2C, Fig 2D, Fig 2F, and Fig 2G, we then extended the equations to simulate the *in vivo* system. Protein degradation was included and was set to be the same for all proteins. Cases with and without HSC70.1 binding to its own mRNA were analysed. For the system without HSC70.1 binding to its own mRNA, the translation rate was adjusted such that the steady state HSC70.1 protein levels were the same as for the case with mRNA binding. *HSC70*.*1* mRNA was assumed to be translated only when free, i.e. not bound to HSC70.1 protein. A stress situation was simulated by a rapid production (at t=200) of mis-folded protein (at a steady-state concentration determined by its production and degradation rate). The binding constant of HSC70.1 to its own mRNA was assumed to be higher than the binding constant of HSC70.1 to misfolded proein. We computed the dynamics of HSC70.1 binding to mis-folded proteins and assumed that this catalyses their refolding. The systems of ordinary differential equations (Table S2) were solved numerically with the LSODA method of the deSolve package (Soetaert *et al*., 2010) in R. The reaction scheme is depicted in the information and all equations and parameters are listed in Table S2.

**Figure 1.**
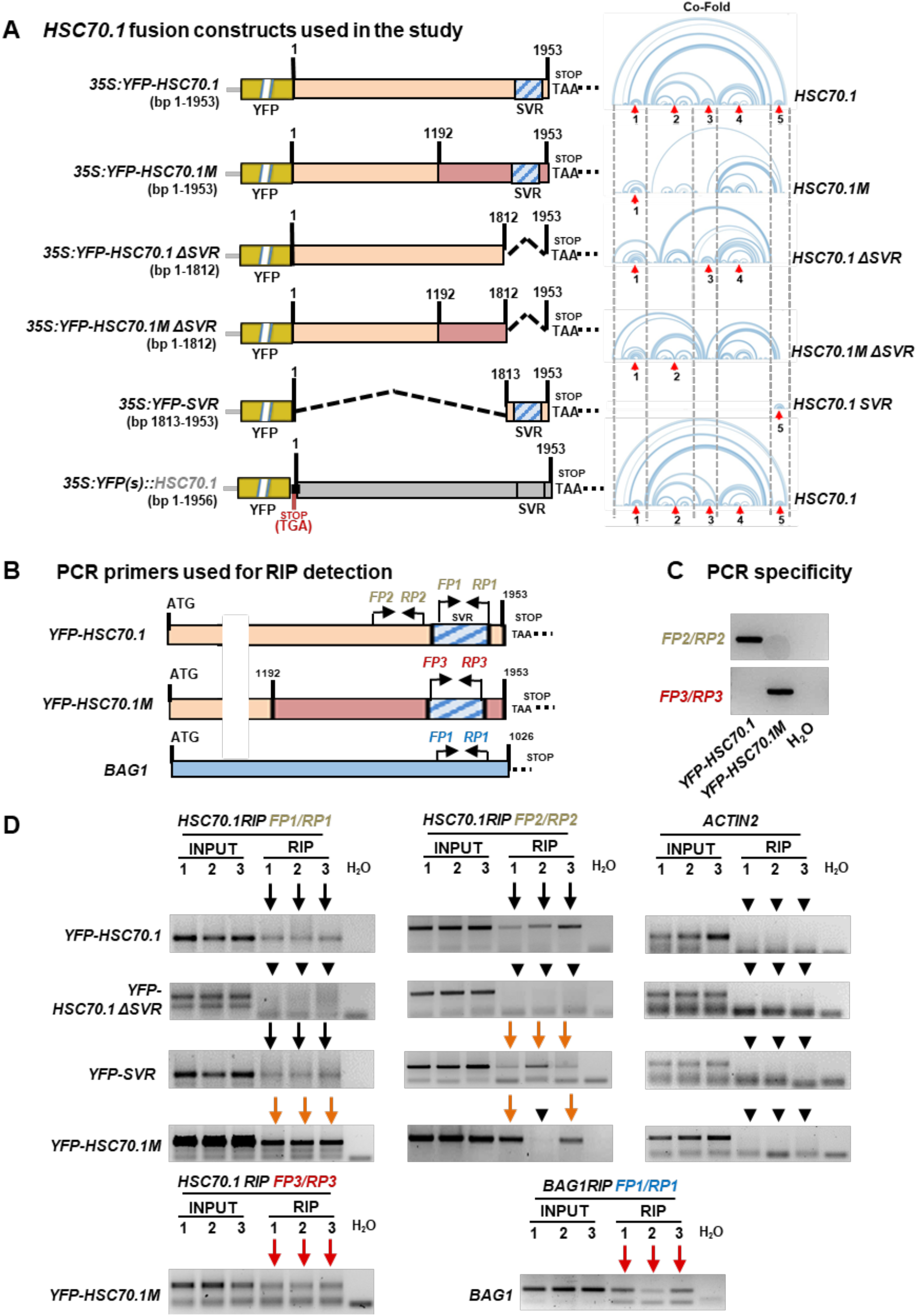
*YFP-HSC70*.*1* fusion constructs and *HSC70*.*1* RNA Immunoprecipitation (RIP) assays. (**A**) Schematic drawing and folding structures of *HSC70*.*1* mRNA *YFP* fusion and deletion constructs used in the study. Left panel: *YFP-HSC70*.*1* (CDS 1-1953); *YFP-HSC70*.*1M* with altered codon usage (wild-type CDS + modified CDS 1192-1953); *YFP-HSC70*.*1 ΔSVR* (CDS 1-1812); *YFP-HSC70*.*1M ΔSVR* (wild-type CDS 1-1191 + modified CDS 1192-1812); *YFP-SVR* (CDS 1813-1953), and *YFP(s)::HSC70*.*1* (wild type CDS 1-1953 preceded by a stop codon). SVR: Short Variable Region. Right panel: Predicted folding of each *HSC70*.*1* construct according to co-transcriptional folding (Co-Fold) (Proctor & Meyer, 2013). Red arrows: five stable regions consistent with RNA folding predicted according to Minimal Free Energy (MFE) (Fig S1 and S2). (**B**) Schematic drawing indicating PCR primers used to detect *HSC70*.*1, HSC70*.*1ΔSVR, HSC70*.*1M*, and *BAG1* transcripts in RIP samples. *HSC70*.*1* FP1/RP1: specific for wild-type (endogenous) *HSC70*.*1* and *YFP-HSC70*.*1*, does not amplify *YFP-HSC70*.*1ΔSVR* and *YFP-HSC70*.*1M. HSC70*.*1* FP2/RP2: specific for wild type *HSC70*.*1, YFP-HSC70*.*1*, and *YFP-HSC70*.*1ΔSVR. HSC70*.*1M* FP3/RP3: specific for *YFP-HSC70*.*1M. BAG1* FP1/RP1: Specifically match to *BAG1* 3’ region. (**C**) Control PCR with FP2/RP2 and FP3/RP3 primer pairs showing that they discriminate between *HSC70*.*1* and *HSC70*.*1M* sequences. (**D**) RT-PCR assays on RNA from *YFP-HSC70*.*1, YFP-HSC70*.*1 ΔSVR, YFP-SVR*, and *YFP-HSC70*.*1M* input and RIP samples. Left and middle panel: RT-PCR assays to detect *HSC70*.*1, HSC70*.*1ΔSVR, SVR, and HSC70*.*1M*. Right panel: control RT-PCR with *ACTIN2* specific primers. Bottom right panel: RT-PCR assays to detect *BAG1* RNA from *YFP-HSC70*.*1* RIP samples. Black arrows: Presence of *HSC70*.*1*. Orange arrows; Presence of endogenous (wild-type) *HSC70*.*1*. Red arrows: Presence of *HSC70*.*1M* and *BAG1* transcript in RIP samples. Arrowheads: No transcript detected. INPUT: DNaseI treated cell extracts used for RIP assays. H_2_O: PCR contamination control with water instead of cDNA.

**Figure 2.**
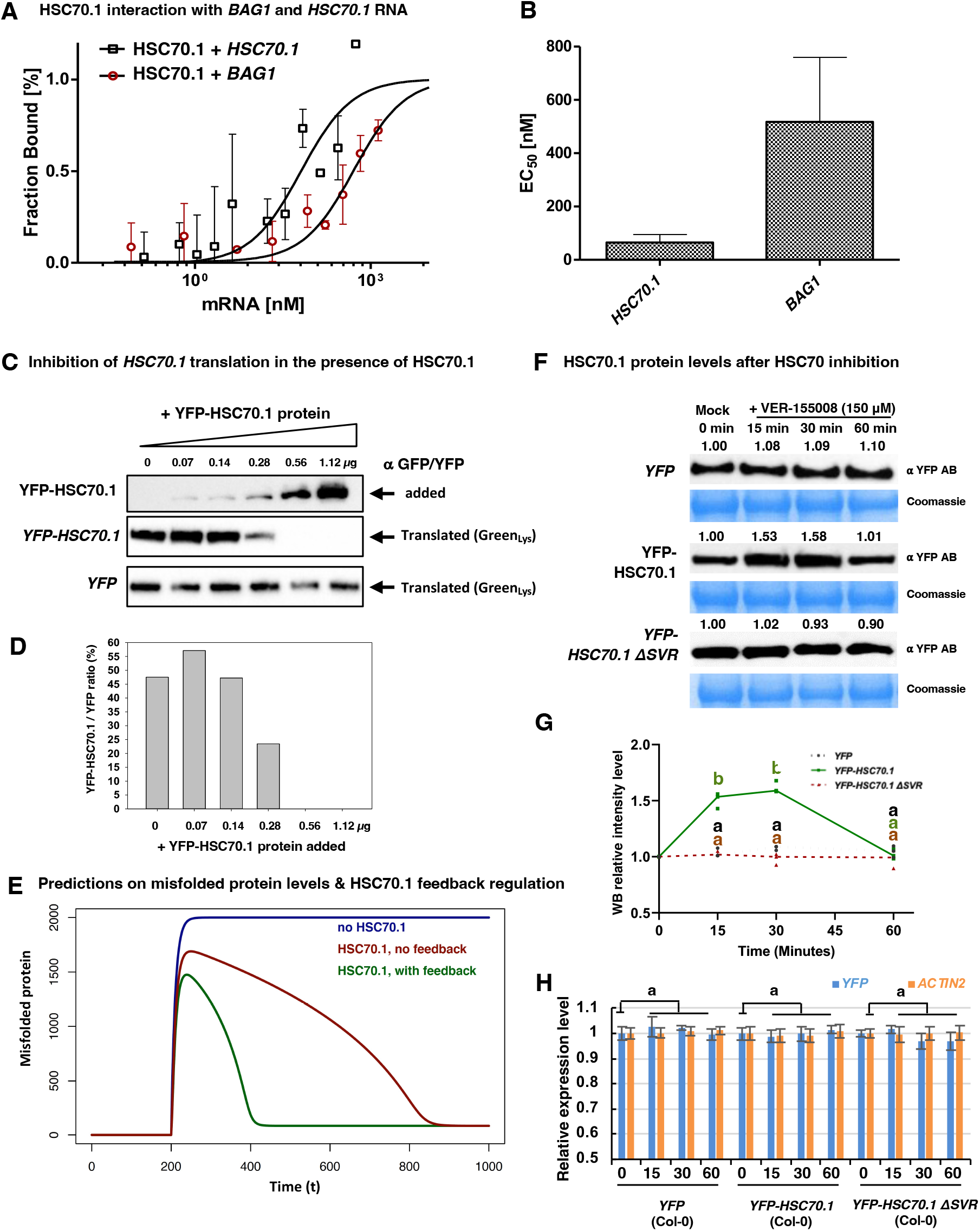
*HSC70*.*1* RNA binding capacity measured by Microscale Thermophoresis and autoregulatory feed-back regulation of translation. (**A**) Microscale Thermophoresis experiment. *In vitro* synthesized *BAG1* and *HSC70*.*1* RNAs ranging between 1 μM and 30 pM were titrated against cell lysates from transgenic plants expressing YFP-HSC70.1 fusion protein. Binding curves were evaluated and plotted as fraction bound against increasing RNA concentrations. Differences in binding are visible as a shift of the sigmoidal binding curve along the x-axis. (**B**) EC50 estimation for both curves. The E_C50_ of *BAG1* towards HSC70.1 is approx. 9 times higher than towards HSC70.1 indicating a higher affinity of HSC70.1 protein towards *HSC70*.*1* compared to *BAG1*. (**C**) Inhibition of *YFP-HSC70*.*1* vs. *YFP* translation in the presence of increasing concentrations of YFP-HSC70.1 protein. YFP-HSC70.1 was extracted from 10 days old transgenic plants and added to the *in vitro* wheat germ expression assay translating either YFP (control) or YFP-HSC70.1. Anti-GFP/YFP antibody was used to detect YFP-HSC70.1 protein addded and Green_lys_ was used to detect newly synthesized YFP-HSC70.1 and YFP. (**D**) Relative ratios of *in vitro* translated YFP-HSC70.1 and YFP protein (see also Fig S6B). (**E**) Model of the predicted effect of HSC70.1 inhibiting its own translation on the refolding of misfolded client proteins. A stress event is assumed to occur (at t=200 s) that gives rise to a sudden increase in misfolded protein. How quickly this amount of misfolded protein deccreases shows how well the HSC70.1 system performs. The model predicts no (blue), slow (red), or a fast (green) refolding of client proteins to due sudden (acute) chaperone demands with no feedback or with translational feedback of HSC70.1 on its own translation (for details see results text, Fig S6C, and Table S3). (**F**) Western Blot (WB) assays on 14 days old YFP, YFP-HSC70.1, and YFP-HSC70.1 ΔSVR transgenic wild-type plants used for HSC70.1 inhibitor (VER-155008) treatment (0, 15 minutes, 30 minutes, and 60 minutes). YFP, YFP-HSC70.1, and YFP-HSC70.1 ΔSVR fusion proteins were detected by YFP antibody. (**G**) Line plot representing average density of bands measured on western blots relative to mock control (n= 3 independent experiments). Significance was calculated using Student’s T-test (two tails); p values indicated by a and b: a,b< 0.001. (**H**) qRT-PCR assays on samples from 14 days old *YFP, YFP-HSC70*.*1*, and *YFP-HSC70*.*1 ΔSVR* transgenic wild-type plants treated with HSC70.1 inhibitor VER-155008 for 0, 15 minutes, 30 minutes, and 60 minutes. Compare with non-treated samples, all treated samples show no significant differences in *YFP* and and *YFP-HSC70*.*1* transcript levels (p values >0.05 ; n = 3 biological replicates; 4 technical replicates). Y axis: relative transcripts levels of *YFP* and *YFP-HSC70*.*1* normalized to *UBQ10*. Error bars: Standard Error.Significance was calculated using Student’s T-test (two tails).

## RESULTS

### HSC70.1 Binds to its Own and Other *HSC70* Transcripts via the C-terminal Short Variable Region (SVR)

Previous studies indicate that human HSC70 chaperones can bind to 3’UTR AU-rich elements of transcripts via their conserved ATPase and substrate binding domains. It was suggested that this interaction stabilizes transcripts found in the animal vasculature (Henics *et al*., 1999; Kishor *et al*., 2013; Kishor *et al*., 2017). Interestingly, the analysis of the whole mammalian RNA interactome indicated the HSC70 C-terminal sequence as a potential RNA binding region (Hentze *et al*., 2018). However, to date, there are no reports on plant HSC70s interacting with mRNAs, and if so, whether their C-terminal SVR sequence is involved in RNA binding. To address these questions we produced transgenic lines expressing YFP fusions of full-length *HSC70*.*1 (YFP-HSC70*.*1)*, deletion variants lacking either the *HSC70*.*1* C-terminal SVR domain (*YFP-HSC70*.*1 ΔSVR*) or the HSC70 substrate and ATPase domain (YFP-SVR), structurally modified *HSC70*.*1* transcripts with alternative codon usage (*YFP-HSC70*.*1M, YFP-HSC70*.*1M ΔSVR)*, and a non-translated *YFP::HSC70*.*1 RNA fusion (YFP(s)::HSC70*.*1*) (Fig 1A; Dataset S1). These *HSC70*.*1* fusions covered also five predicted stable *HSC70*.*1* RNA structures as indicated by co-transcriptional folding (Co-Fold) (Proctor & Meyer, 2013) and minimal free energy RNA folding (MFE) algorithms (Zuker & Stiegler, 1981; Lorenz *et al*., 2011) (Fig 1A, Fig S1 and S2). We confirmed also that these various Y*FP-HSC70*.*1* fusion constructs were stably expressed in *A. thaliana* and that the intracellular cytosolic and nuclear protein localization was not altered by the introduced changes (Fig S3 and S4).

To identify RNAs that bind to HSC70.1, we performed RNA-protein immunoprecipitation (RIP) assays on extracts from *35S::YFP-HSC70*.*1* and *35S::erYFP* (control) transgenic plants using anti-YFP/GFP-trap beads. RIP RNA samples (three biological replicates per line) were submitted to RNAseq (paired-end reads) and analyzed for significant transcript enrichment (log2>=1, p<=0.05 FDR) using the *A. thaliana* Araport 11 cDNA database. According to this analysis 13 RNAs were found to be significantly enriched in the YFP-HSC70.1 RIP samples compared with erYFP control samples (Fig S5). Notably, 9 of the 13 enriched transcripts encode RNA binding proteins and 4 of the 13 transcripts had previously been identified as being potentially graft mobile (Fig S5) (Thieme *et al*., 2015). The most significant (FDR p-value 6.10E-4) enriched HSC70.1 interacting RNA sequence was the 5’ region of a transcript encoding the highly conserved HSC70 co-chaperone BAG1 (Fig S5) known to bind to and regulate HSP70 chaperone activity (Lee *et al*., 2016). Interestingly, also the *HSC70*.*1* transcript was found significantly enriched (FDR p-value 0.05) pointing towards a potential autoregulatory function based on RNA translation.

To substantiate this finding, we performed additional RNA co-immunoprecipitation (RIP) assays on samples from *35S::YFP-HSC70*.*1, 35S::YFP-HSC70*.*1 ΔSVR, 35S::YFP-SVR* and *35S::YFP-HSC70*.*1M* transgenic plants (Fig 1B-D). These assays, combined with RT-PCR assays to detect the presence of *HSC70*.*1*, indicated that *HSC70*.*1* mRNA binds to YFP-HSC70.1 and YFP-SVR but not to YFP-HSC70.1 ΔSVR (Fig 1D), suggesting that the SVR region binds the *HSC70* transcript. Notably, the endogenously produced wild-type *HSC70*.*1* transcript (Fig 1D) could be detected in RT-PCR assays performed on YFP-HSC70.1M RIP samples using a set of specific PCR primers discriminating between *HSC70*.*1M* and wild-type *HSC70*.*1*, suggesting the interaction with mRNA can occur also after it has been translated. Also, *BAG1* RNA interaction with HSC70.1 protein was confirmed by RT-PCR assays on YFP-HSC70.1 RIP samples (Fig 1D). To further corroborate *in planta* that HSC70.1 binds to the identified RNAs, we performed microscale thermophoresis experiments using cell extracts of transgenic plants expressing YFP-HSC70.1 or YFP alone (control). These extracts were supplemented with *in vitro* produced tagged *HSC70*.*1* and *BAG1* mRNA and thermophoretic mobility was measured (Fig 2A and B). These assays established that HSC70.1 protein binds to *HSC70*.*1* and *BAG1* RNA in the nanomolar range. Thus, our data indicate that similar to the human HSP70 (Zimmer *et al*., 2001) that the *A. thaliana* HSC70.1 chaperone binds its own mRNA via its C-terminal SVR sequence.

### HSC70.1 Protein Binding its mRNA Negatively Regulates Its Own Translation

Through the use of transcriptional and translational inhibitors, previous studies with Drosophila cells found evidence that HSP70 protein translation can be controlled both transcriptionally and post-transcriptionally by regulation of both *HSP70* mRNA synthesis and mRNA destabilization and that this regulatory pathway may depend on the abundance of its protein (DiDomenico *et al*., 1982). In line with this prediction, we found that Arabidopsis HSC70.1 binds to its own transcript.

To further substantiate that HSC70 chaperones might regulate directly its own translation, we first tested whether increasing concentrations of YFP-HSC70.1 protein impacts the degree of translation of *YFP-HSC70*.*1* transcript in *in vitro* assays. Note that YFP-HSC70.1 translation could only be detected *in vitro* when relatively high amounts of bivalent Mg2+ was added, which is known to alter and stabilize RNA structures (for details see Methods). We next enriched YFP-HSC70.1 protein from transgenic seedlings using specific YFP AB beads and added these in increasing concentrations to the *in vitro* translation system and assayed its effect on translation (Fig 2C and D, and Fig S6A and B). In the presence of > 0.5 μg of YFP-HSC70.1, very low or no translation of YFP-HSC70.1 was detected. This was in stark contrast to YFP translation where no inhibition was observed in the presence of HSC70 protein (Fig 2C and D, and Fig S6B). Notably, with low amounts of added YFP-HSC70.1, we detected a slight initial increase of translation (Fig 2C and D, and Fig S6B). This was expected as HSC70.1 chaperones are known to accelerate ribosomal activity.

Negative or balancing feedback is known to provide a means of maintaining a desired level (Maxwell, 1868) but the implications of this for HSC70.1 were not clear. To explore how the negative feedback regulation of HSC70.1 might impact its ability to act as a chaperone responding to stress, we built a computational model of the system based on the above observations. There are some key assumptions built into the model, some of which require further validation. 1) We assumed that the HSC70.1 binds to its own transcript with a defined equilibrium binding constant, 2) *HSC70*.*1* transcript cannot be translated when bound to HSC70.1 protein, 3) HSC70.1 protein has a lower equilibrium binding constant to misfolded proteins than to its own transcript, i.e. in the presence of misfolded protein, the complex of HSC70.1 protein with *HSC70*.*1* transcript will dissociate and binding to misfolded protein will dominate over HSC70.1 protein binding to its own transcript. Thus, when no or low concentrations misfolded proteins are present, HSC70.1 will be at a defined level, determined by the binding constant to its own transcript, and most *HSC70*.*1* mRNA will be in this bound, non-translatable form. We computed what happens when the amount of misfolded protein is increased (Fig 2E and Fig S6C). Using the above assumptions, the model predicts that with a translation-regulating feedback loop, the response time for achieving the same chaperone availability is shorter relative to an equivalent system without feedback (Fig 2E), consistent with other biological systems (Rosenfeld *et al*., 2002).

To validate these predictions from the model, we conducted an *in vivo* experiment in which we measured the abundance of constitutively expressed YFP (control), YFP-HSC70.1, and YFP-HSC70.1 ΔSVR (control, not binding its own transcript) in 14 days old seedlings after applying the HSP70s specific inhibitor VER-155008 (Fig 2F and G). VER-155008 is an adenosine analog targeting specifically the HSC70s ATPase binding domain impairing HSC70 chaperone function (Williamson *et al*., 2009; Schlecht *et al*., 2013; Merret *et al*., 2015). We expected that the inhibitor limits HSC70 substrate turnover, leading to an acute chaperone demand. YFP-HSC70.1 bound to its own transcript maintains a reservoir of both *HSC70*.*1* mRNA and protein, which can be made available on demand. Thus, a system with such as negative feedback due to chaperone-mRNA binding should show increased translation once mRNA is released from its interaction, whereas the two YFP-HSC70.1 ΔSVR and YFP proteins not binding to their own transcripts, should exhibit a much slower change of translation as HSC70.1 would first need to be transcribed. In other words, in the presence of VER-155008 an acute response would be that more HSC70 protein is produced independently of gene transcription. As anticipated only YFP-HSC70.1 and not YFP or YFP-HSC70.1 ΔSVR protein levels increased significantly by approx. 1.5x after 15 and 30 minutes of VER-155008 incubation (Fig 2F and G, and Table S1). Notably, YFP-HSC70.1 protein levels returned to normal after 1 hour showing no significant differences to preincubation levels. Also transcription from the constitutive *35S* promoter was not significantly changed by VER-155008 treatment as indicated by qRT-PCR assays (Fig 2H). Thus, the detected specific and significant increase of YFP-HSC70.1 protein in such a short period was due to an increased *YFP-HSC70*.*1* translation and not due to a change in gene expression of the *35S* promoter constructs.

Taken together, our data suggest that *A. thaliana* HSC70.1 interacts with its own mRNA and therby regulates its own translation. This constitutes a negative regulatory feedback loop between *HSC70*.*1* transcript translation and cellular HSC70.1 protein demands and constitutes a stable/homeostatic system that can respond rapidly to fluctuating environmental conditions (Fig 2E and Fig S6C).

### *HSC70*.*1* Transcript Mobility Depends on Specific mRNA Sequences Encoding the SVR RNA-binding Motif

Full-length *A. thaliana HSC70*.*1* transcript lacking its endogenous 5’ and 3’ UTR sequences fused to *GFP* moves from above-ground tissue (shoot, scion) to roots (root-stock) in hypocotyl grafted plants (Yang *et al*., 2019). To analyze whether transport of *HSC70*.*1* transcript depends on specific mRNA sequences, we first grafted the *YFP-HSC70*.*1* transgenic lines (TG) with wild type (Col-0) and assayed the presence of HSC70.1 fusion constructs in heterologous tissues in plants grown in liquid culture 30 days after grafting (Fig 3A and Fig S7). Consistent with previous results (Yang *et al*., 2019), confocal laser scanning microscopy (CLSM) and RT-PCR assays revealed that both full-length HSC70.1 fusion protein and *HSC70*.*1* fusion transcript produced in transgenic wild-type shoots could be detected in wild-type roots (Fig 3A and B).

**Figure 3.**
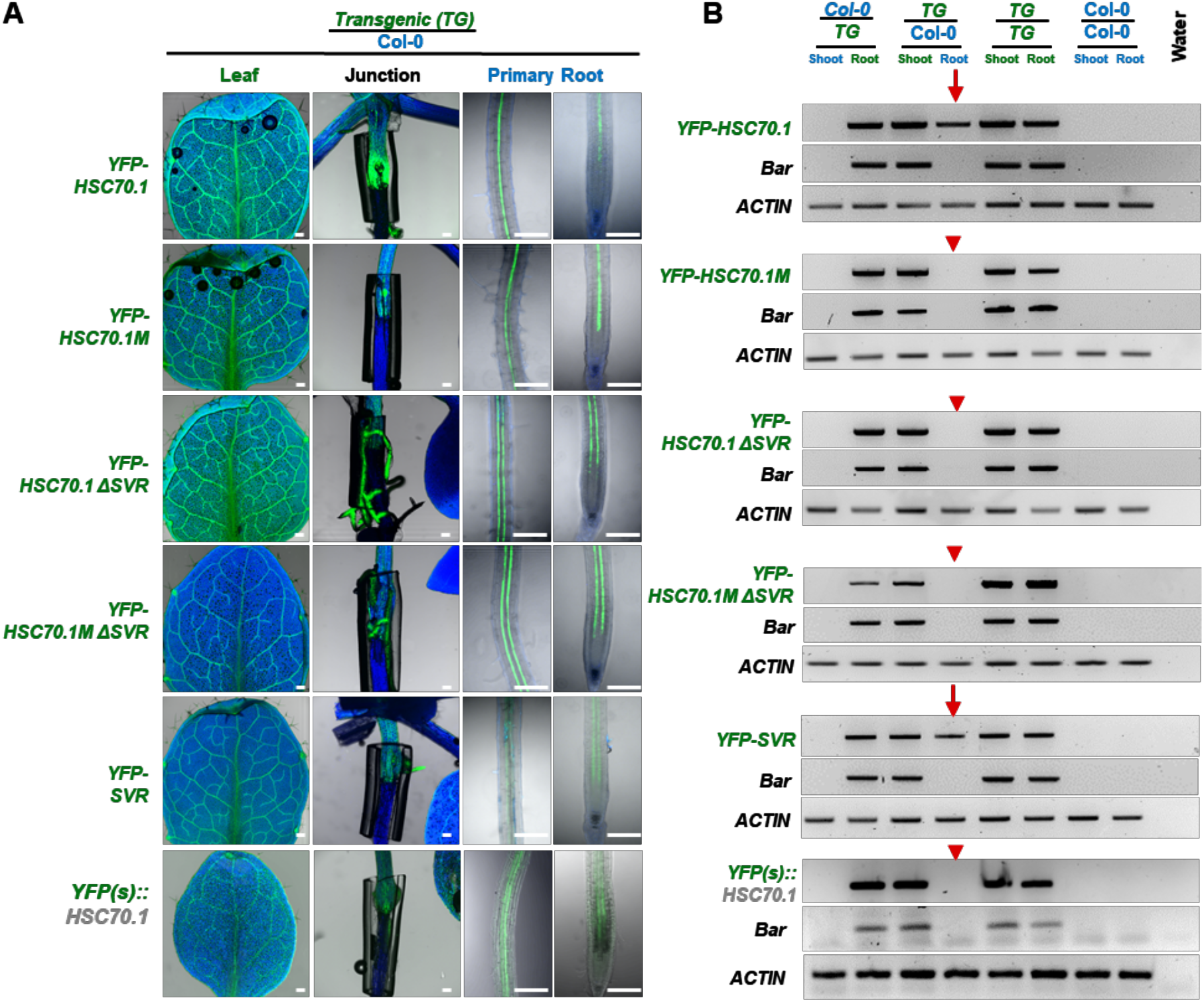
Mobility of the *YFP-HSC70*.*1* transcript fusion variants. (**A**) YFP fluorescence detected by CLSM in leaves, graft junctions, and primary roots of grafted *YFP-* fusion transgenic / wild-type (Col-0) plants. Blue color: auto-fluorescent. Green color: YFP fluorescence. Three independent lines (each n>30 plants) were used for each graft combination showing a similar YFP signal distribution. Bar: 200 μm. (**B**) RT-PCR detection of *YFP* fusion constructs in root and shoot samples from grafted plants. Grafted plant material was pooled (n>6) and tested for transcript presence in shoots and roots (45 PCR cycles). Note that transcript absence was additionally confirmed by 50 PCR cycles. Red arrows: presence, red arrowheads: absence of *YFP RNA* fusion constructs in grafted wild type (Col-0) tissue.

We next tested the mobility of the *HSC70*.*1M* construct which has a changed codon usage in the 3’ half of the *HSC70*.*1* protein*-*coding sequence. This variant *HSC70*.*1M* produces a wild-type HSC70 protein but shows changes in the predicted folding of 4 out of the 5 predicted stable RNA folding structures (Fig 1A and Fig S1; Dataset S1). In grafted plants, fluorescent YFP-HSC70.1M fusion protein produced in scions was present abundantly in wild-type roots (Fig 3A); however, in the wild-type roots of the same grafted plants, a *YFP-HSC70*.*1M* transcript was not detectable by highly sensitive RT-PCR assays (Fig 3B). This suggests that HSC70.1 protein mobility is independent of *HSC70*.*1* RNA mobility and that *HSC70*.*1* transcript transport depends on its RNA structure enclosing the C-terminal half of the encoding sequence including the SVR region (Fig 1A, and Fig S1 and S2).

To address whether the *HSC70*.*1* transcript sequence encoding the RNA-binding SVR region is sufficient and necessary in mediating *HSC70*.*1* transcript mobility, we examined the mobility of truncated *YFP-HSC70*.*1* constructs lacking the *SVR* RNA sequence *(YFP-HSC70*.*1 ΔSVR and YFP-HSC70*.*1M ΔSVR*; lacking the 3’ SVR region stretching from base +1813 to +1953). Both SVR deletion transcripts were not detected in grafted wild-type roots in RT-PCR assays (Fig 3B), indicating this mRNA region is necessary for *HSC70*.*1* transcript mobility. We next tested whether the identified SVR encoding RNA sequence is sufficient to mediate mobility by grafting *YFP-SVR* expressing transgenic shoots with wild-type roots (Fig 3A). Here, the transcript was detected in grafted wild-type roots suggesting that the fused *SVR* transcript sequence mediates *YFP* RNA transport (Fig 3B).

We next asked whether graft-mobility of the *YFP-HSC70*.*1* transcript depends on its translation. For this we introduced a stop codon between the YFP and HSC70.1 RNA fusion. This construct does not produce an HSC70.1 fusion protein in transgenic plants but contains the SVR encoding RNA sequences mediating *HSC70*.*1* RNA mobility (Fig 1A and Fig S4). Notably, this fusion construct was not sufficient to mediate transport of the fused *YFP* RNA (Fig 3B), suggesting that translation of the encoded HSC70.1 protein is required to mediate transport of its own mRNA.

These combined results indicate that the 140 bases of the SVR sequence of *HSC70*.*1* contains an RNA motif that is sufficient and necessary for transcript mobility and that its transport depends on the presence of HSC70.1 protein produced by the same transcript.

### Arabidopsis HSC70.1 Protein Transport is Independent of its SVR Motif

In contrast to the reported role of the pumpkin CmHSC70.1 SVR amino acid sequence in mediating intercellular mobility in microinjection assays (Aoki *et al*., 2002), the *A. thaliana* HSC70.1 SVR protein sequence does not seem to be necessary for HSC70.1 shoot-to-root transport in grafted plants (Fig 3A). Previous work indicated that free YFP protein diffuses via the phloem vasculature to the root in grafted plants (Yang *et al*., 2019) and that proteins > 60 kDa cannot freely move between cells and via the phloem to grafted roots (Paultre *et al*., 2016). In line, compared to YFP, which is unloaded into the root apical cells, the distribution of the YFP-HSC70.1 fusions appeared to be restricted to the root vasculature (Fig 3A and Fig S8A). Also, there is no obvious difference in the cellular distribution of graft-mobile YFP-HSC70.1 fusions moving to wild-type roots. All YFP-HSC70.1 protein fusion variants were found in the grafted root vasculature independent of the SVR region and of their sizes which ranged from > 100 kDa (YFP-HSC70.1) to ∼36 kDa (YFP-SVR) (Fig S3 and S8*A*). In particular the large YFP fusion proteins produced by *YFP-HSC70*.*1 ΔSVR, YFP-HSC70*.*1M ΔSVR*, and *YFP-HSC70*.*1M* transgenic plants, whose transcripts were not detected in the grafted wild-type roots (Fig 3B) appeared as full-length protein fusions in wild-type roots (Fig S8*B*) indicading that they are not delivered as smaller degradation products which can diffuse similar to free YFP. Thus, it seems that the HSC70.1 SVR domain does not play a crucial role in providing long-distant transport of HSC70.1 protein in grafted *A. thaliana* plants.

To address whether the *A. thaliana* HSC70.1 SVR sequence plays a similar role in local cell-to-cell transport as seen with pumpkin HSC70.1 (Aoki *et al*., 2002), we performed transient single-cell expression assays after biolistic gold particle bombardment of *Nicotiana benthamiana* and *Nicotiana sylvestris* leaves. Previous work revealed that free GFP with a size of ∼ 27 kDa expressed in a single leaf epidermal cell diffuses via plasmodesmata to neighboring cells. In a similar line, we used free YFP protein expressed from the binary vector used to produce the YFP-HSC70.1 fusion construct as a reference for local cell-to-cell transport. In control assays with YFP (∼31 kDa) 19% (8 out of 42) of bombarded cells showed YFP fluorescence in one neighboring cell and 9.5% (4 out of 42) in more than one neighboring cell 36h after bombardment. In contrast to YFP alone, YFP-HSC70.1 (∼102 kDa), YFP-HSC70.1 ΔSVR (∼98 kDa), and YFP-SVR (∼36 kDa) appeared in 27.3% (27 out of 99), 37.3% (21 out of 68) and 30.9% (40 out of 107), in 7.1% (7 out of 99), 16.2% (11 out of 68) and 28% (30 out of 107) in more than one neighboring cell, respectively (Fig 4A and B). The intercellular mobility of the YFP fusion proteins seemed to be independent of cell size (Fig 4C). However, cell-to-cell mobility of the HSC70.1 fusion proteins was significantly higher than YFP free diffusion (Fig 4). Here, in contrast to previous reports on pumpkin HSC70 mobility assays based on microinjection (Aoki *et al*., 2002), the Arabidopsis HSC70.1 SVR domain - although its presence significantly increased YFP mobility compared to YFP alone (Fig 4) - seems not to be essential for local HSC70.1 cell-to-cell movement.

**Figure 4.**
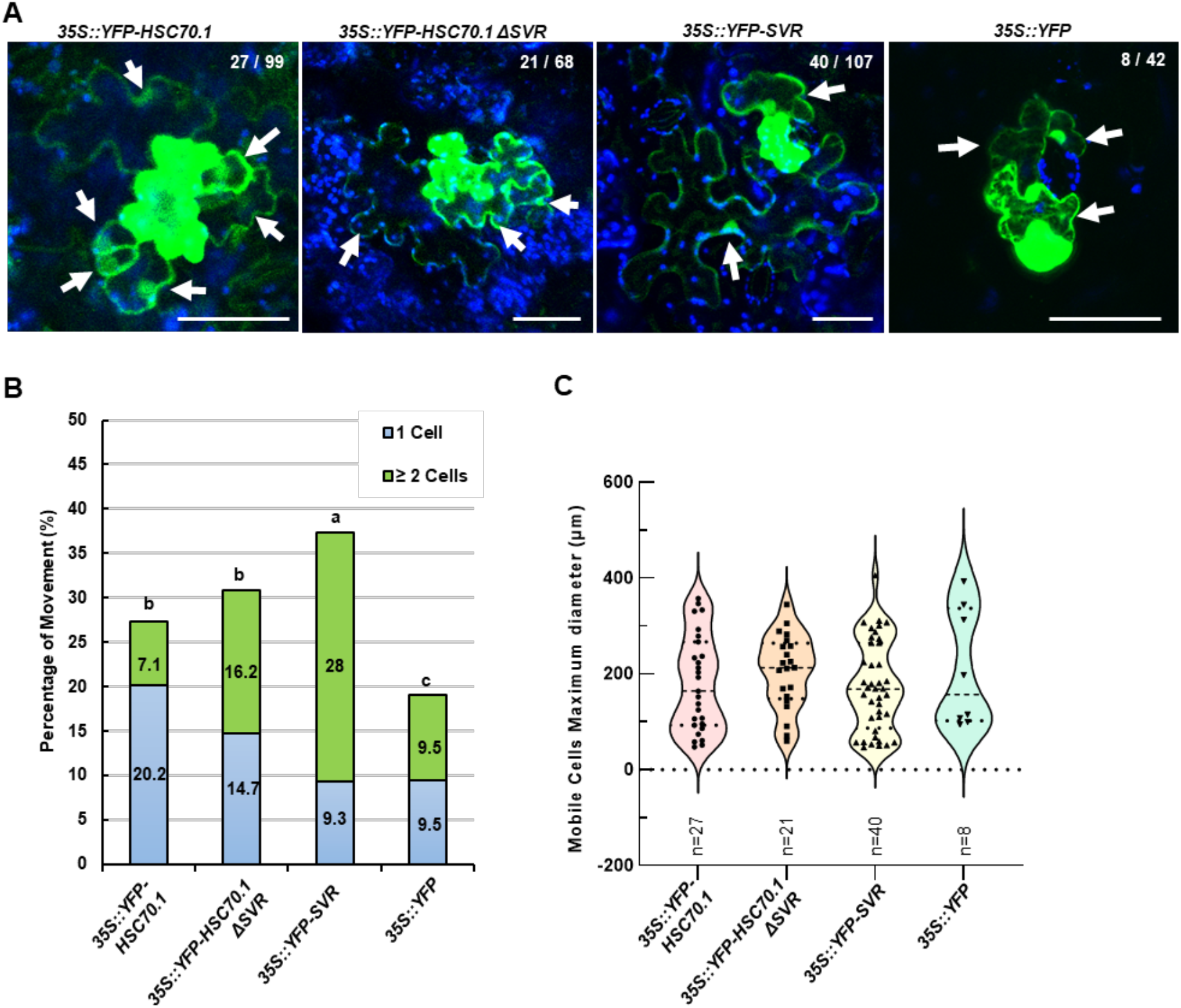
Single-cell expression and intercellular movement of YFP-HSP70.1 fusions. (**A**) Representative CLSM images *N. benthamiana* epidermal cells bombarded with *35S* promoter constructs expressing YFP-HSC70.1, YFP-HSC70.1ΔSVR, YFP-SVR, or YFP. Green: YFP fluorescence. Blue: chloroplastic auto-fluorescence. White arrows: neighboring cells with YFP fluorescence. Numbers indicate the fraction of analyzed YFP expressing cells that show a YFP signal in neighboring cells. Bar: 100 μm. (**B**) Intercellular movement ratio in % of all expressing cells and fraction showing movement to one or two and more (?2) neighboring cells. Significance was calculated using Student’s T-test (two tails); p values indicated by a, b and c: a,b< 0.01; b,c< 0.05; a,c<0.001. (**C**) Size distribution of cells with detected incidents of cell-to-cell mobility. Note that the maximum diameter was measured and that individual points indicate cells in which the YFP construct moved.

### Mobile *YFP-HSC70*.*1* Transcript Complements the Root Growth of *hsc70*.*1 hsp70*.*4* mutants

10 out of 18 *A. thaliana HSP70* family members were found to be graft-mobile in heterografted plants (Thieme *et al*., 2015) (Table S2), indicating that there may be a benefit of *HSC70s* acting non-cell-autonomously. No noticeable phenotypes were reported for *hsc70* single mutants, however *hsc70*.*1 hsp70*.*4* double and *hsp70*.*2 hsp70*.*4 hsp70*.*5* triple mutants develop strong phenotypes with curly/round leaves, twisted petioles, thin stems, early flowering, and short siliques (Leng *et al*., 2017). Thus, to test whether mobility of *HSC70*.*1* protein or transcript is necessary for normal Arabidopsis growth and development, we produced independent *hsc70*.*1 hsp70*.*4* double mutant lines (n>59) expressing mobile *YFP-HSC70*.*1* and non-mobile *YFP-HSC70*.*1M* transcripts (Fig S9). Note that, in contrast to transgenic *YFP-HSC70*.*1* and *YFP-HSC70*.*1M* wild-type lines, the transgenic *YFP-HSC70*.*1* and *YFP-HSC70*.*1M hsc70*.*1 hsp70*.*4* double mutants showed a high degree of *YFP-HSC70* gene-silencing (Fig S3A-E compared to Fig S9A-E). Relatively high *HSC70*.*1* silencing was also apparent in *hsc70*.*1* single mutants and may be explained by strong up-regulation of other HSC70 family members in these mutants (Fig S9G) (Taipale *et al*., 2014; Leng *et al*., 2017). Despite this, we were able to select relatively stable *YFP-HSC70*.*1* (n=2) and *YFP-HSC70*.*1M* (n=3) transgenic lines allowing us to ask whether these fusion constructs can complement the aberrant *hsc70*.*1 hsp70*.*4* growth phenotypes. Expression of the constructs in transgenic lines was confirmed by western blot and quantitative RT PCR assays, revealing that the transformed mutant lines - with the exception of *YFP-HSC70*.*1 hsc70*.*1 hsp70*.*4* transgenic line #*2 –* exhibit no obvious differences in HSC70 fusion protein expression levels and intracellular distribution of the fusion construct (Fig S9F and G). Also, no significant difference in germination time and rate between the transgenic and wild-type lines was noticed in the used lines (Fig S10).

We next measured primary root growth of the selected plant lines (Fig 5). This revealed that wild type, *hsc70*.*1*, and *hsp70*.*4* single mutants have significantly longer primary roots than *hsc70*.*1 hsp70*.*4* double mutants. The two independent *YFP-HSC70*.*1 #1 (hsc70*.*1 hsp70*.*4*) and *YFP-HSC70*.*1 #2 (hsc70*.*1 hsp70*.*4*) transgenic lines showed similar primary root length as wild-type plants. In contrast, the three independent *hsc70*.*1 hsp70*.*4* transgenic plant lines expressing the non-mobile *YFP-HSC70*.*1M (*line *#*6, #9, and #11) transcript showed no significant root growth differences compared to *hsc70*.*1 hsp70*.*4* double mutants. These results imply that long-distance movement of *HSC70*.*1* mRNA is necessary to rescue delayed root growth in *hsc70*.*1 hsp70*.*4* double mutants. Notably, although the YFP-HSC70.1 protein produced by the non-mobile *YFP-HSC70*.*1M* transcript was still transported to wild-type roots (Fig 3A and Fig S8), it failed to rescue primary root growth (Fig 5) which suggests that the mobile HSC70.1 transcript and not the HSC70.1 mobile protein is required to restore root growth.

**Figure 5.**
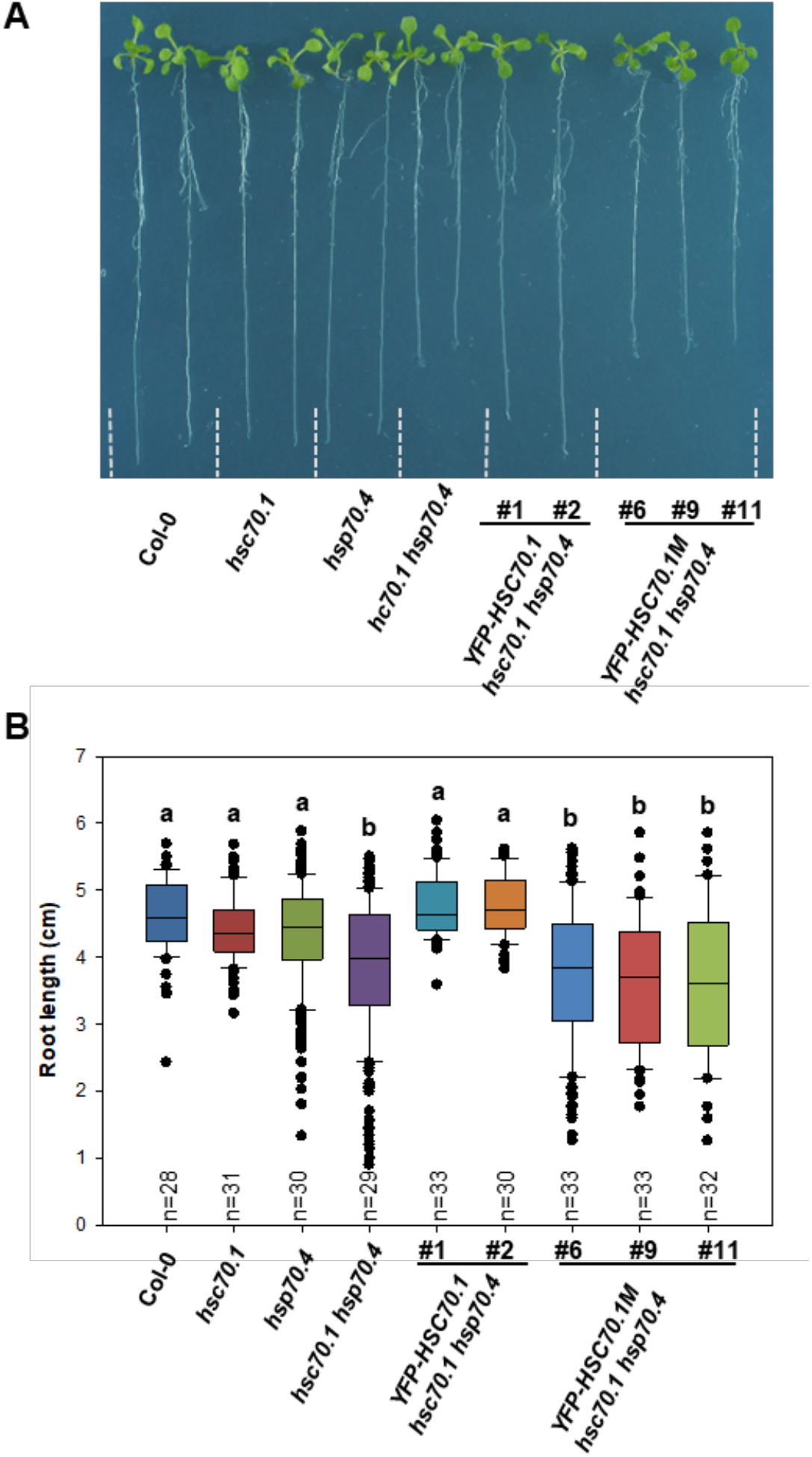
Mobile *HSC70*.*1* transcript rescues *hsc70*.*1 hsp70*.*4* root growth. (**A**) Representative pictures of analyzed wild-type (Col-0), *hsc70*.*1, hsp70*.*4, hsc70*.*1 hsp70*.*4, HSC70*.*1 #1* (*hsc70*.*1 hsp70*.*4), HSC70*.*1 #2* (*hsc70*.*1 hsp70*.*4*), *HSC70*.*1M #6* (*hsc70*.*1 hsp70*.*4*), *HSC70*.*1M #9* (*hsc70*.*1 hsp70*.*4*) and *HSC70*.*1M-1 #11* (*hsc70*.*1 hsp70*.*4*) plants 14 days after germination. (**B**) Quantitative data of measured primary root length of wild-type and indicated mutant plants. Box plot graph: Boxes denote variation between datasets and means; n = number of analyzed plants; error bar: ± SE; black dots: measurements out of ± SE range. Significance was calculated using Student’s T-test (two tails); p value indicated by a and b: a, b<0.001.

### Mobile *HSC70*.*1* Transcript Rescues Early Flowering of *hsc70*.*1 hsp70*.*4* Double Mutants

In our phenotypic analysis of *hsc70*.*1 hsp70*.*4* double mutants, we noticed an early flowering (bolting) phenotype under long-day growth conditions. Thus, we compared the flowering time of wild type, *hsc70*.*1* and *hsp70*.*4* single mutants, *hsc70*.*1 hsp70*.*4* double mutants, and transgenic *YFP-HSC70*.*1* and *YFP-HSC70*.*1M hsc70*.*1 hsp70*.*4* double mutants (Fig 6). This comparison revealed that wild type, *hsc70*.*1* and *hsp70*.*4* single mutants, and *YFP-HSC70*.*1* (*hsc70*.*1 hsp70*.*4*) transgenics bolted 32 - 36 days after germination (DAG), whereas *hsc70*.*1 hsp70*.*4* double mutants and *YFP-HSC70*.*1M* (*hsc70*.*1 hsp70*.*4*) transgenics bolted significantly earlier between 27 - 32 DAG and 27 – 34 DAG, respectively (Fig 6B). To address whether early bolting was due to growth delay we counted the number of rosette leaves present on flowering plants. At time of bolting the wild type, *hsc70*.*1* and *hsp70*.*4* single mutants had 15 to 20 rosette leaves, which was significantly more than detected on *hsc70*.*1 hsp70*.*4* double mutant plants with 13 to 17 rosette leaves (Fig 6C). *YFP-HSC70*.*1 expressing hsc70*.*1 hsp70*.*4* double mutants formed 14 to 21 rosette leaves which was similar to wild type and *hsc70*.*1* and *hsp70*.*4* single mutants. In contrast, the three independent *YFP-HSC70*.*1M* (*hsc70*.*1 hsp70*.*4*) transgenic plants showed 11 to 18 rosette leaves which was not significantly different compared to *hsc70*.*1 hsp70*.*4* double mutants (Fig 6C).

**Figure 6.**
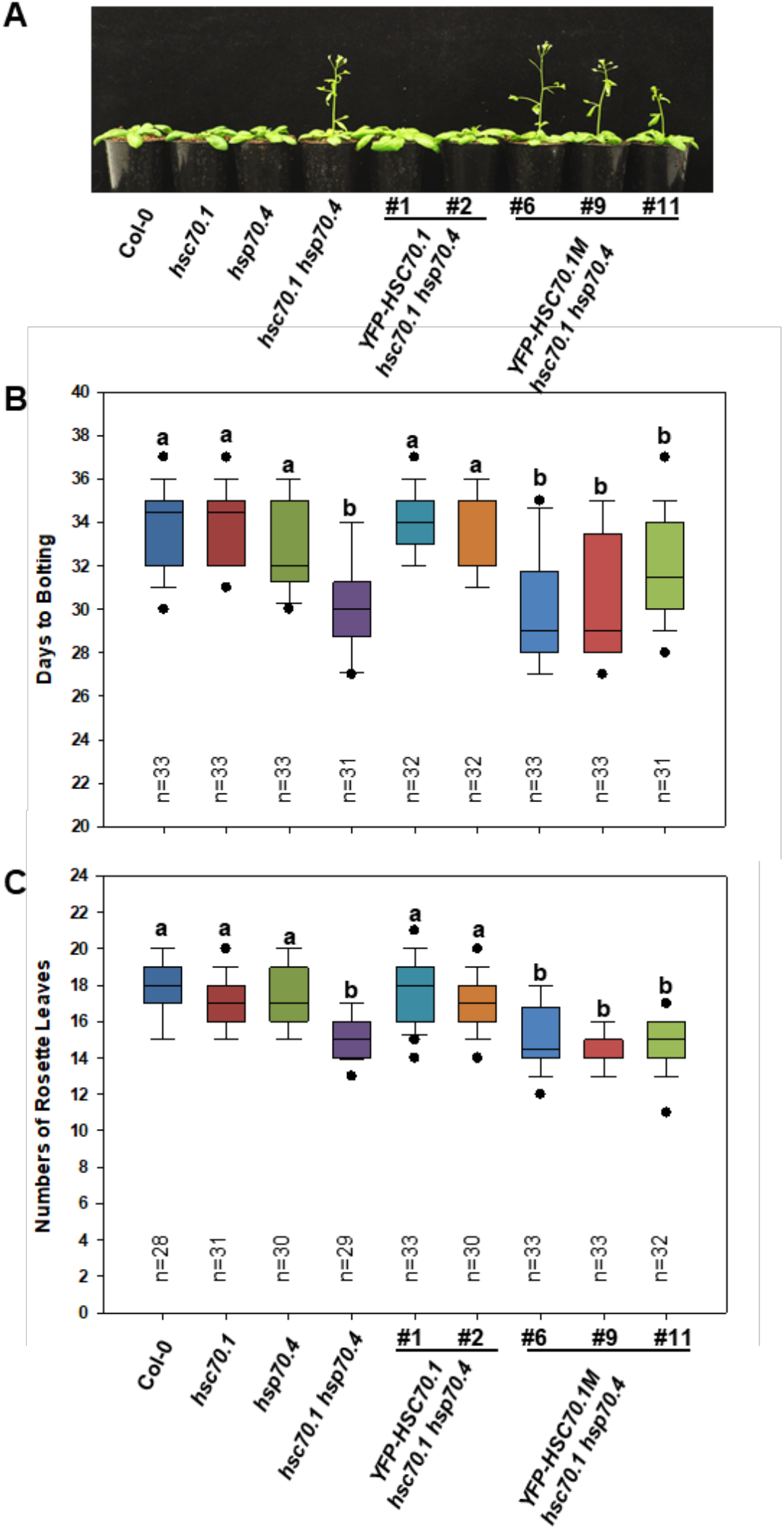
Mobile *HSC70*.*1* transcript rescues *hsc70*.*1 hsp70*.*4* early flowering phenotype. (**A**) Representative pictures of analyzed wild-type (Col-0), *hsc70*.*1, hsp70*.*4, hsc70*.*1 hsp70*.*4, HSC70*.*1 #1* (*hsc70*.*1 hsp70*.*4), HSC70*.*1 #2* (*hsc70*.*1 hsp70*.*4*), *HSC70*.*1M #6* (*hsc70*.*1 hsp70*.*4*), *HSC70*.*1M #9* (*hsc70*.*1 hsp70*.*4*) and *HSC70*.*1M-1 #11* (*hsc70*.*1 hsp70*.*4*) plants 30 days after germination. (**B**) Age of plants at bolding of wild-type and indicated mutant plants. (**C**) Numbers of rosette leaves at time of bolding of wild-type and indicated mutant plants. Box plot graph: Boxes indicate variation between datasets and means; n = number of plants analyzed; error bar: ± SE; black dots: measurements out of range ± SE; significance was calculated using Student’s T-test (two tails); p value indicated by a and b: a, b<0.001.

These results indicate that mobile HSC70.1 protein alone could not rescue the root growth and early flowering phenotype of *hsc70*.*1 hsp70*.*4* double mutants (Fig 5 and Fig 6). As mentioned above, the *HSC70*.*1M* transcript was not graft-mobile, but its translated protein was moving between cells in single cell expression assays and in grafted plants. Thus, we asked whether the *YFP(s)::HSC70*.*1* fusion transcript not producing an HSC70.1 protein (Fig 1A and Fig S4) and not showing mobility in grafted plants (Fig 3B) is also not sufficient to complement the *hsc70*.*1 hsp70*.*4* mutant flowering phenotype. For this purpose we used *hsc70*.*1 hsp70*.*4* double mutant transgenic lines (expressing the *YFP(s)::HSC70*.*1* fusion variant (Fig S4) and compared these to wild-type (Col-0) transgenic (n>13) and *hsc70*.*1 hsp70*.*4* mutants. Three independent, stable *YFP(s)::HSC70*.*1 (hsc70*.*1 hsp70*.*4)* transgenic lines were selected and their root growth and flowering time were measured (Fig S11 and Fig S12). All three *YFP(s)::HSC70*.*1* transgenic lines showed no significant root growth and flowering time differences compared to *hsc70*.*1 hsp70*.*4* double mutant plants. This indicates that non-translated *HSC70*.*1* RNA fused to YFP was not sufficient to rescue the delayed root growth and early flowering.

To evaluate whether phenotype complementation might depend on the ubiquitius expression of the used *35S* promoter-driven *HSC70*.*1* fusion construct, we grafted *hsp70*.*4* single with *hsc70*.*1 hsp70*.*4* double mutant plants (Fig 7). Here, homografted *hsp70*.*4* and homografted *hsc70*.*1 hsp70*.*4* plants served as a control for the flowering time phenotype. Homografted *hsc70*.*1 hsp70*.*4* / *hsc70*.*1 hsp70*.*4* plants started to bolt after 27 to 30 days whereas homografted *hsp70*.*4* / *hsp70*.*4* plants started to bolt significantly later after 31 to 34 days. In contrast to homografted *hsc70*.*1 hsp70*.*4* / *hsc70*.*1 hsp70*.*4* plants, *hsc70*.*1 hsp70*.*4* / *hsp70*.*4* heterografted mutants started to bolt significantly later after 31 to 34 days (Fig 7A and C) suggesting that *hsp70*.*4* root-stocks could suppress the earlier *hsc70*.*1 hsp70*.*4* scion flowering phenotype. Notably, the complementation of the early flowering phenotype correlated with *HSC70*.*1* mRNA movement from *hsp70*.*4* rootstock to *hsc70*.*1 hsp70*.*4* scion in flowering plants as revealed by RT PCR assays (Fig 7B) and which was not detected in young plants (Fig 3B). In summary, the complementation and grafting assays suggest that both, *HSC70*.*1* transcript mobility and HSC70.1 protein expression, are necessary to promote primary root growth and suppress early flowering of *hsc70*.*1 hsp70*.*4* mutants. Furthermore, this suggests that, that *HSC70*.1 transcript mobility - although *HSC70*.*1* is expressed in all tissues - is necessary for proper plant growth.

**Figure 7.** Grafted *hsp70*.*4* mutants producing wild-type *HSC70*.*1* transcript complements early *hsc70*.*1 hsp70*.*4* flowering. (**A**) Representative pictures of root/shoot grafted *hsc70*.*1 hsp70*.*4* mutant plants 29 days after grafting. (**B**) RT-PCR assays confirming presence of mobile *HSC70*.*1* transcript produced in *hsp70*.*4* mutant tissue in heterologous *hsc70*.*1 hsp70*.*4* root and shoot tissue (arrowheads). (**C**) Time of flowering (bolding) in days after grafting. Box plots: Boxes indicate the variation between datasets and means; 16 grafted plants were analyzed for each graft combination; error bar: ± SE; black dots: measurements out of range ± SE; significance was calculated using Student’s T-test (two tails); p-value indicated by a and b: a, b<0.001.

## DISCUSSION

The Arabidopsis HSC70.1 is a housekeeping chaperone and as such is involved in many important pathways related to plant growth (Vierling, 1991; Sung *et al*., 2001; Cazale *et al*., 2009; Clement *et al*., 2011; Leng *et al*., 2017). HSC70.1 is expressed to equally high levels in all plant cell types and approx. 2-3 times higher in dividing and endreduplicating cells (Apelt *et al*., 2022). Although it was reported to be induced by severe heat stress, the transcription of *HSC70*.*1* does not appear to be changed by a first mild and a later applied severe heat stress (Olas *et al*., 2021). In line, there is a low correlation between gene expression and protein abundance of HSC70 family members suggesting that a post-transcriptional control mechanism is in place regulating HSP70s activity (Berka *et al*., 2022). HSC70 chaperones are *bona fide* RNA binding proteins proposed to stabilize binding transcripts (Kishor *et al*., 2013; Kishor *et al*., 2017) yet the relevance of this in plants and of HSC70 RNA binding capacity has remained unclear.

In animal cells, translation of HSC70 has been predicted to be regulated on the translational level, depending on its abundance (DiDomenico *et al*., 1982), although a direct interaction between HSC70 protein and its transcript was not established in these assays. Our thermophoretic RNA-binding studies and translational assays with plant produced HSC70 samples (Fig 2) show that HSC70.1 binds to its own transcript with high affinity and that abundant HSC70 protein this inhibits its own translation. In line, specific inhibition of HSC70 chaperone activity in plants results in a specific increase of HSC70 translation after 15 minutes independent of transcriptional changes (Fig 2). Taken together this points to a negative feedback regulation of HSC70.1 that inhibits its own translation (Fig 2E). High demand for chaperone activity would dissociate HSC70.1 from its own transcript, freeing up HSC70.1 protein and making mRNA available for translation.

Given the redundancy with >18 HSC70 chaperones expressed in *A. thaliana*, the highly conserved sequence, the housekeeping function of chaperones in facilitating folding of client proteins, and the suggested role of chaperones in compensating random somatic mutations (Queitsch *et al*., 2002), it might be surprising that 10 of these HSC70-related transcripts were identified as mobile (Thieme *et al*., 2015). Interestingly, the rescue of early flowering and wild-type-like root growth was provided by mobile *HSC70* and not by non-mobile transcript versions (Fig 5 and 6). Mobile HSC70.1 protein alone, whose mobility does not seem to depend on *HSC70*.*1* transcript mobility, was not sufficient to compensate for loss of HSC70 function in receiving cells. This is not surprising given the high concentrations of HSC70 that are required in cells and the low detected transport rates of protein in grafted plants.

Another question was whether *HSC70*.*1* transcript mobility depends on the capacity of HSC70.1 protein to bind its RNA. We could demonstrate that *HSC70*.*1* RNA transport over graft junctions depends on a defined RNA sequence motif that coincides with the SVR encoding region mediating HSC70 protein binding to its RNA. Thus, it seems that both features, the capacity of the SVR motif to bind to RNA (Fig 1*D*) and the presence SVR encoding sequence, are essential for *HSC70*.*1* transcript transport (Fig 3). However, although the SVR motif significantly enhances intercellular protein transport (Fig 4) its presence seems not to be essential for long-distance transport in grafted plants (Fig 3). Notably, a YFP expressing transcript fused with non-translated *HSC70*.*1* RNA was not graft mobile (Fig 1*C*), which suggests that the *SVR* RNA sequence has to be translated to be recognized as an RNA transport motif. Here, a simple explanation could be that *HSC70*.*1* mRNAs form with their own encoded HSC70.1 protein graft-mobile ribonuleoprotein complexes at ribosomes during their translation process when not acutely needed for folding its own or other nascent proteins.

In summary, we established that the HSC70.1 C-terminal SVR motif is needed for long-distance transport, enhances intercellular protein transport and binds to *HSC70*.*1* and other mobile transcripts. Furthermore, the HSC70.1 chaperone seems to negatively regulate its own translation and that the interaction with its own transcript is needed to be transported as mRNA and protein between cells and tissues. One could speculate that these features have the potential to create a non-cell autonomous homestatic chaperone system allowing cells/tissues to adopt to acute local stresses within minutes and to balance chaperone activity over longer time periods between distant tissues (Fig 8). We hypothesize that the autoregulatory feedback regulation of HSC70 translation acts locally within minutes whereas the slow long-distance transport of HSC70 RNA is required for coordinated growth of distant tissues when parts of a plant are locally exposed to stresses over extended time periods and under normal conditions. This model would be consistent with our finding that mobile *HSC70*.1 transcipt, and not the mobile HSC70.1 protein, is the critical determinant for normal growth (Fig 5 and 7). This model is also based on the observation that transported mRNA is translated in receiving cells (Zhang, W *et al*., 2016). Given that HSC70 chaperones are also non-cell-autonomous proteins in animals (De Maio, 2014), we propose that both intercellular transport of HSC70s and negative feedback regulation of its own translation is an evolutionarily conserved feature potentially constituting a relatively simple way to coordinate and maintain chaperone homeostasis within and between tissues.

**Figure 8.**
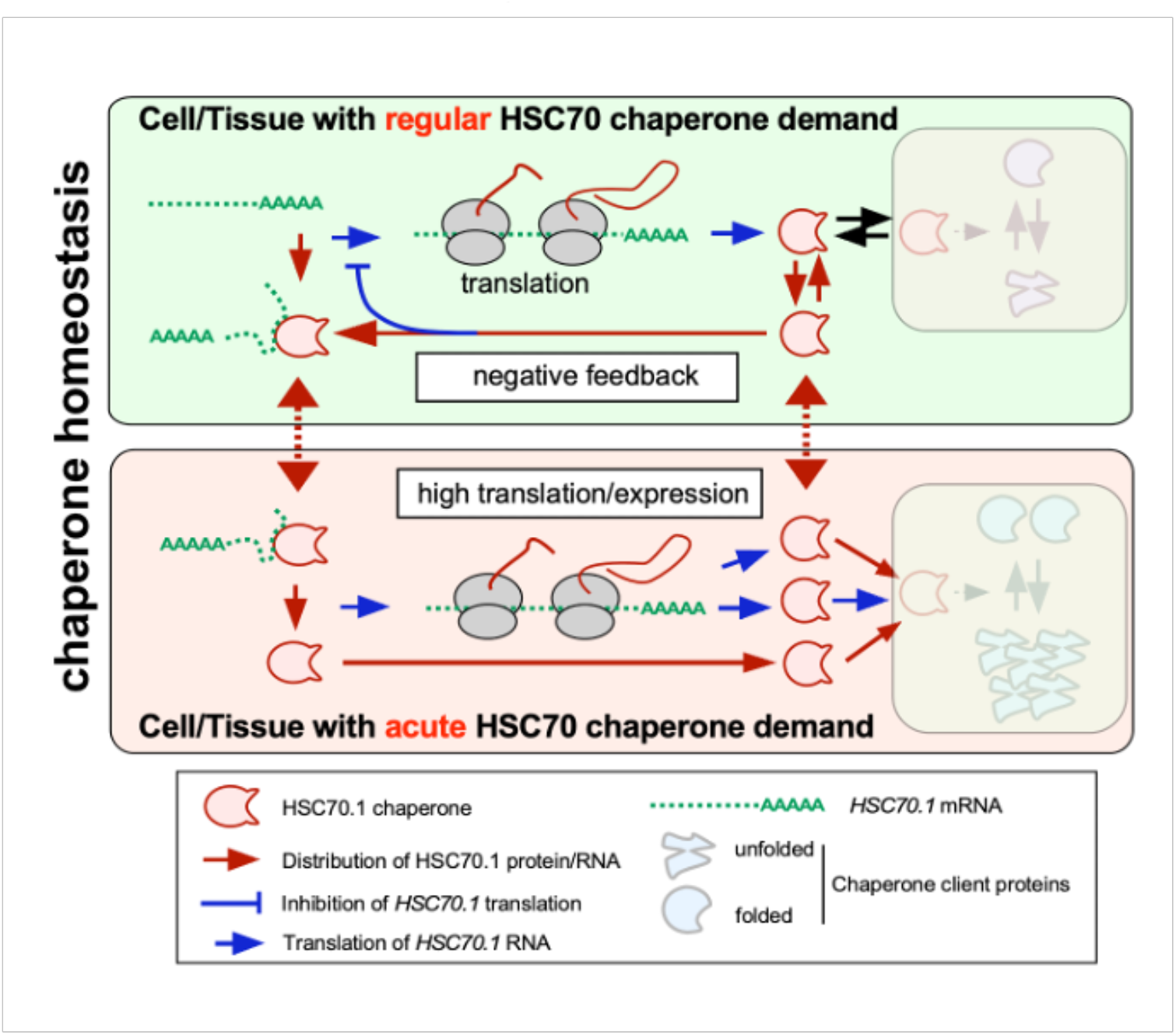
Speculative model of HSC70 functioning as a non-cell-autonomous mobile chaperone regulating its own translation. HSC70 can block its own translation by binding its mRNA (negative feedback). In non-stressed cells (with regular chaperone activity demants) this will result in a balanzed pool of complexes of HSC70 bound to its own mRNA and of translated HSC70 protein available for folding of client proteins. HSC70 transcript and protein, possibly as HSC70 transcript-protein complexes, can move between cells and to distant tissues. The function of this movement remains to be determined, but potentially plays a role in maintaining homeostatic chaperone levels between tissues/cells over longer time periods. Under stress conditions, the HSC70 transcript-protein complexes may dissociate, freeing up both active HSC70 protein and releaving the auto-inhibitory activity on its own translation, thus allowing for a rapid (minutes) response to sudden increases in chaperone demand. After longer time periods (hours) HSC70 gene expression can be induced in a stressed tissue/cell producing more of HSC70 protein and RNA. As HSC70 transcript and protein can move they would be delivered to non-stressed tissues/cells. In recipient cells imported additional HSC70 protein might interfere with translation and lower chaperone activity in distant tissues resulting in coordinated decreased growth in e.g. a locally stressed plant. In parallel, recipient cells/tissues migh be able to adopt faster to anticipated stress by additionally received *HSC70* mRNA. Both features, HSC70 protein - transcript auto-inhibitory activity and transcript transport between cells, may enable a multicellular organism to establish chaperone homeostasis between e.g. stressed and non-stressed tissues over long time periods providing additional robustness in stress conditions to ensure coordinated growth within and between tissues.

## AVAILABILITY

All study data are included in the article and/or SI Appendix Figs and Datasets. The RIP RNAseq data are available for download at the European Nucleotide Archive (ENA) under the accession number PRJEB44090 (https://www.ebi.ac.uk/ena).

## SUPPLEMENTARY DATA

Supplementary Data are available at NAR online.

## AUTHOR CONTRIBUTION

L.Y and F.K. conceived the project and suggested experiments; L.Y produced the transgenic plants, performed the grafting experiments, analyzed the growth and mobility data, and with Y.Z. performed bombardment experiments; SF. W. with L.Y performed the in vitro and in vivo expression experiments. Y.X. performed RNA IP experiments. F.K. analyzed the RIP RNAseq data; R.J.M. developed the model for the simulation of feed-back regulation; M.T. analyzed the *HSC70*.*1* RNA sequences; S.O. with J.K. suggested and S.O. performed the thermophoretic RNA interaction assays. F.K. assisted by L.Y. wrote the manuscript. All authors read and contributed to the manuscript text and expressed their consent to submit the final version of the manuscript.

## ACKNOWLEDGEMENT

We thank Saurabh Gupta (MPI-MPP, Golm, Germany) for his advice to analyze the RIP deep sequencing data and Dana Schindelasch (MPI-MPP, Golm, Germany) for her excellent technical support; Wei Su (Fu Dan University, Shanghai, China) for providing *hsc70*.*1 hsp70*.*4* double mutant seeds and the anonymous reviewers of the initial version of the manuscript for their helpful comments and corrections.

## FUNDING

This work was supported by MPI-MPP internal funds to F.K.; L.Y., Y. Z., and Y. X. were supported by a Chinese Scholarship Council (CSC) PhD Stipend; This article is part of a project that has received funding from the European Research Council (ERC) under the European Union’s Horizon 2020 research and innovation programme (Grant agreement No. 810131).

## CONFLICT OF INTEREST

The authors declare no conflict of interests.

## Notes

### Competing Interest Statement

The authors have declared no competing interest.

