## Supplementary material for "Non-cell-autonomous HSC70.1 chaperone displays homeostatic feed-back regulation by binding its own mRNA": Dataset

### Supplementary Dataset S1

#### Sequences of the *YFP-HSC70.1* constructs used in the study.

Green colour background indicates *35S* promoter; Yellow colour background indicates *YFP*; Light blue colour background indicates *HSC70.1* (open reading frame CDS) RNA sequences; Red colour background indicates the deleted *HSC70.1* sequences; Dark red colour background indicates the HSC70.1 modified sequence. Gray colour background indicates *OCS* terminator sequences. **Bold:** start/stop codon.

***1. 35S::YFP:HSC70.1::OCS***

TGAGACTTTTCAACAAAGGATAATTTCGGGAAACCTCCTCGGATTCCATTGCCCAGCTATCTGTCACTTCATCGAAAGGACAGTAGAAAAGGAA
GGTGGCTCCTACAAATGCCATCATTGCGATAAAGGAAAGGCTATCATTCAAGATCTCTCTGCCGACAGTGGTCCCAAAGATGGACCCCCACCCA
CGAGGAGCATCGTGGAAAAAGAAGACGTTCCAACCACGTCTTCAAAGCAAGTGGATTGATGTGACATCTCCACTGACGTAAGGGATGACGCACA
ATCCCACTATCCTTCGCAAGACCCTTCCTCTATATAAGGAAGTTCATTTCATTTGGAGAGGACACGCTCGAGTATAAGAGCTCTATTTTTACAA
CAATTACCAACAACAACAAACAACAAACAACATTACAATTACATTTACAATTACC **ATG** GTG AGC AAG GGC GAG GAG CTG TTC ACC
GGG GTG GTG CCC ATC CTG GTC GAG CTG GAC GGC GAC GTA AAC GGC CAC AAG TTC AGC GTG TCC GGC GAG GGC
GAG GGC GAT GCC ACC TAC GGC AAG CTG ACC CTG AAG TTC ATC TGC ACC ACC GGC AAG CTG CCC GTG CCC TGG
CCC ACC CTC GTG ACC ACC TTC GGC TAC GGC CTG CAG TGC TTC GCC CGC TAC CCC GAC CAC ATG AAG CAG CAC
GAC TTC TTC AAG TCC GCC ATG CCC GAA GGC TAC GTC CAG GAG CGC ACC ATC TTC TTC AAG GAC GAC GGC AAC
TAC AAG ACC CGC GCC GAG GTG AAG TTC GAG GGC GAC ACC CTG GTG AAC CGC ATC GAG CTG AAG GGC ATC GAC
TTC AAG GAG GAC GGC AAC ATC CTG GGG CAC AAG CTG GAG TAC AAC TAC AAC AGC CAC AAC GTC TAT ATC ATG
GCC GAC AAG CAG AAG AAC GGC ATC AAG GTG AAC TTC AAG ATC CGC CAC AAC ATC GAG GAC GGC AGC GTG CAG
CTC GCC GAC CAC TAC CAG CAG AAC ACC CCC ATC GGC GAC GGC CCC GTG CTG CTG CCC GAC AAC CAC TAC CTG
AGC TAC CAG TCC GCC CTG AGC AAA GAC CCC AAC GAG AAG CGC GAT CAC ATG GTC CTG CTG GAG TTC GTG ACC
GCC GCC GGG ATC ACT CTC GGC ATG GAC GAG CTG TAC AAG TCC GGA CTC AGA TCT CGA GCT CAA GCT TCG AAT
TCT GCA GTC GAC GGT ACC GCG GGC CCG GGA TCA TCA ACA AGT Ttg tac aaa aaa gca ggc tcc acc atg gga
acc aat tca gtc gac tgg atc cgc ATG TCG GGT AAA GGA GAA GGA CCA GCT ATC GGT ATC GAT CTT GGT ACC
ACT TAC TCT TGC GTC GGA GTA TGG CAA CAC GAC CGT GTT GAG ATC ATT GCT AAT GAT CAA GGA AAC AGA ACC
ACG CCA TCT TAC GTT GCT TTC ACC GAC TCC GAG AGG TTG ATC GGT GAC GCA GCT AAG AAT CAG GTC GCC ATG
AAC CCC GTT AAC ACC GTT TTC GAC GCT AAG AGG TTG ATC GGT CGT CGT TTC TCT GAC AGC TCT GTT CAG AGT
GAC ATG AAA TTG TGG CCA TTC AAG ATT CAA GCC GGA CCT GCC GAT AAG CCA ATG ATC TAC GTC GAA TAC
AAG GGT GAA GAG AAA GAG TTC GCA GCT GAG GAG ATT TCT TCC ATG GTT CTT ATT AAG ATG CGT GAG ATT GCT
GAG GCT TAC CTT GGT GTC ACA ATC AAG AAC GCC GTT GTT ACC GTT CCA GCT TAC TTC AAC GAC TCT CAG CGT
CAG GCT ACA AAG GAT GCT GGT GTC ATC GCT GGT TTG AAC GTT ATG CGA ATC ATC AAC GAG CCT ACA GCC GCC
GCT ATT GCC TAC GGT CTT GAC AAA AAG GCT ACC AGC GTT GGA GAG AAG AAT GTT CTT ATC TTC GAT CTT GGT
GGT GGC ACT TTT GAT GTC TCT CTT CTT ACC ATT GAA GAG GGT ATC TTT GAG GTG AAG GCA ACT GCT GGT GAC
ACC CAT CTT GGT GGG GAA GAT TTT GAC AAC AGA ATG GTT AAC CAC TTT GTC CAA GAG TTC AAG AGG AAG AGT
AAG AAG GAT ATC ACC GGT AAC CCA AGA GCT CTT AGG AGG TTG AGA ACT TCC TGT GAG AGA GCG AAG AGG ACT
CTT TCT TCC ACT GCT CAG ACC ACC ATC GAG ATT GAC TCT CTA TAC GAG GGT ATC GAC TTC TAC TCC ACC ATC
ACC CGT GCT AGA TTT GAG GAG CTC AAC ATG GAT CTC TTC AGG AAG TGT ATG GAG CCA GTT GAG AAG TGT CTT
CGT GAT GCT AAG ATG GAC AAG AGC ACT GTT CAT GAT GTT GTC CTT GTT GGT GGT TCT ACC CGT ATC CCT AAG
GTT CAG CAA TTG CTC CAG GAC TTC TTC AAC GGC AAA GAG CTT TGC AAG TCT ATT AAC CCT GAT GAG GCT GTT
GCC TAC GGT GCT GCT GTC CAG GGA GCT ATT CTC AGC GGT GAA GGA AAC GAG AAG GTT CAA GAT CTT CTA TTG
CTC GAT GTC ACT CCT CTC TCC CTT GGT TTG GAA ACT GCC GGT GGT GTC ATG ACC ACT TTG ATC CCA AGG AAC
ACA ACC ATC CCA ACC AAG AAG GAA CAA GTC TTC TCC ACC TAC TCA GAC AAC CAA CCC GGT GTG TTG ATC CAG
GTG TAC GAA GGA GAG AGA GCC AGA ACC AAG GAC AAC AAC CTT CTT GGT AAA TTT GAG CTC TCC GGA ATT CCT
CCA GCT CCT CGT GGT GTC CCC CAG ATC ACA GTC TGC TTT GAC ATT GAT GCC AAT GGT ATC CTC AAT GTC TCT
GCT GAG GAC AAG ACC ACC GGA CAG AAG AAC AAG ATC ACC ATC ACC AAT GAC AAG GGT CGT CTC TCC AAG GAT
GAG ATT GAG AAG ATG GTT CAA GAG GCT GAG AAG TAC AAG TCC GAA GAC GAG GAG CAC AAG AAG AAG GTT GAA
GCC AAG AAC GCT CTC GAG AAC TAC GCT TAC AAC ATG AGG AAC ACC ATC CAA GAC GAG AAG ATT GGT GAG AAG
CTC CCG GCT GCA GAC AAG AAG AAG ATC GAG GAT TCT ATT GAG CAG GCG ATT CAA TGG CTC GAG GGT AAC CAG
TTG GCT GAG GCT GAT GAG TTC GAA GAC AAG ATG AAG GAA TTG GAG AGC ATC TGC AAC CCA ATC ATT GCC AAG
ATG TAC CAA GGA GCT GGT GGT GAA GCC GGT GGT CCA GGT GCC TCT GGT ATG GAC GAT GAT GCT CCC CCT GCT
TCA GGC GGT GCT GGA CCT AAG ATC GAG GAG GTC GAC **TAA**
cggccgcactcgagatatctagacccagctttCTTGTACAAAGTGGTGCTAGGTGAGTCT
AGAGAGTTAATTAAGACCCGGGACTAGTCCCTAGAGTCCTGCTTTAATGAGATATGCGAGACGCCTATGATCGCATGATATTTGCTTTCAATTC
TGTTGTGCACGTTGTAAAAAACCTGAGCATGTGTAGCTCAGATCCTTACCGCCGGTTTCGGTTCATTCTAATGAATATATCACCCGTTACTATC
GTATTTTTATGAATAATATTCTCCGTTCAATTTACTGATTGTACCCTACTACTTATATGTACAATATTAAAATGAAAACAATATATTGTGCTGA
ATAGGTTTATAGCGACATCTATGATAGAGCGCCACAATAACAAACAATTGCGTTTTATTATTACAAATCCAATTTTAAAAAAAGCGGCAGAACC
GGTCAAACCTAAAAGACTGATTACATAAATCTTATTCAAATTTCAAAAGTGCCCCAGGGGCTAGTATCTACGACACACCGAGCGGCGAACTAAT
AACGCTCACTGAAGGGAACTCCGGTTCCCCGCCGGCGCGCATGGGTGAGATTCCTTGAAGTTGAGTATTGGCCGTCCGCTCTACCGAAAGTTAC
GGGCACCATTCAACCCGGTCCAGCACGGCGGCCGGGTAACCGACTTGCTGCCCCGAGAATTATGCAGCATTTTTTTGGTGTATGTGGGCCCCAA
ATGAAGTGCAGGTCAAACCTTGACAGTGACGACAAATCGTTGGGCGGGTCCAGGGCGAATTTTGCGACAACATGTCGAGGCTCAG

***2. 35S::YFP:HSC70.1 ΔSVR (1-1812)::OCS***

TGAGACTTT......ACAATTACC **ATG** GTG AGC AAG...... CTG TAC AAG TCC GGA CTC AGA TCT CGA GCT CAA GCT
TCG AAT TCT GCA GTC GAC GGT ACC GCG GGC CCG GGA TCA TCA ACA AGT Ttg tac aaa aaa gca ggc tcc acc
atg gga acc aat tca gtc gac tgg atc cgc ATG TCG GGT AAA GGA GAA GGA CCA GCT ATC GGT ATC GAT CTT
GGT ACC ACT TAC TCT TGC GTC GGA GTA TGG CAA CAC GAC CGT GTT GAG ATC ATT GCT AAT GAT CAA GGA AAC
AGA ACC ACG CCA TCT TAC GTT GCT TTC ACC GAC TCC GAG AGG TTG ATC GGT GAC GCA GCT AAG AAT CAG GTC
GCC ATG AAC CCC GTT AAC ACC GTT TTC GAC GCT AAG AGG TTG ATC GGT CGT CGT TTC TCT GAC AGC TCT GTT
CAG AGT GAC ATG AAA TTG TGG CCA TTC AAG ATT CAA GCC GGA CCT GCC GAT AAG CCA ATG ATC TAC GTC
GAA TAC AAG GGT GAA GAG AAA GAG TTC GCA GCT GAG GAG ATT TCT TCC ATG GTT CTT ATT AAG ATG CGT GAG

ATT GCT GAG GCT TAC CTT GGT GTC ACA ATC AAG AAC GCC GTT GTT ACC GTT CCA GCT TAC TTC AAC GAC TCT
CAG CGT CAG GCT ACA AAG GAT GCT GGT GTC ATC GCT GGT TTG AAC GTT ATG CGA ATC ATC AAC GAG CCT ACA
GCC GCC GCT ATT GCC TAC GGT CTT GAC AAA AAG GCT ACC AGC GTT GGA GAG AAG AAT GTT CTT ATC TTC GAT
CTT GGT GGT GGC ACT TTT GAT GTC TCT CTT CTT ACC ATT GAA GAG GGT ATC TTT GAG GTG AAG GCA ACT GCT
GGT GAC ACC CAT CTT GGT GGG GAA GAT TTT GAC AAC AGA ATG GTT AAC CAC TTT GTC CAA GAG TTC AAG AGG
AAG AGT AAG AAG GAT ATC ACC GGT AAC CCA AGA GCT CTT AGG AGG TTG AGA ACT TCC TGT GAG AGA GCG AAG
AGG ACT CTT TCT TCC ACT GCT CAG ACC ACC ATC GAG ATT GAC TCT CTA TAC GAG GGT ATC GAC TTC TAC TCC
ACC ATC ACC CGT GCT AGA TTT GAG GAG CTC AAC ATG GAT CTC TTC AGG AAG TGT ATG GAG CCA GTT GAG AAG
TGT CTT CGT GAT GCT AAG ATG GAC AAG AGC ACT GTT CAT GAT GTT GTC CTT GTT GGT GGT TCT ACC CGT ATC
CCT AAG GTT CAG CAA TTG CTC CAG GAC TTC TTC AAC GGC AAA GAG CTT TGC AAG TCT ATT AAC CCT GAT GAG
GCT GTT GCC TAC GGT GCT GCT GTC CAG GGA GCT ATT CTC AGC GGT GAA GGA AAC GAG AAG GTT CAA GAT CTT
CTA TTG CTC GAT GTC ACT CCT CTC TCC CTT GGT TTG GAA ACT GCC GGT GGT GTC ATG ACC ACT TTG ATC CCA
AGG AAC ACA ACC ATC CCA ACC AAG AAG GAA CAA GTC TTC TCC ACC TAC TCA GAC AAC CAA CCC GGT GTG TTG
ATC CAG GTG TAC GAA GGA GAG AGA GCC AGA ACC AAG GAC AAC AAC CTT CTT GGT AAA TTT GAG CTC TCC GGA
ATT CCT CCA GCT CCT CGT GGT GTC CCC CAG ATC ACA GTC TGC TTT GAC ATT GAT GCC AAT GGT ATC CTC AAT
GTC TCT GCT GAG GAC AAG ACC ACC GGA CAG AAG AAC AAG ATC ACC ATC ACC AAT GAC AAG GGT CGT CTC TCC
AAG GAT GAG ATT GAG AAG ATG GTT CAA GAG GCT GAG AAG TAC AAG TCC GAA GAC GAG GAG CAC AAG AAG AAG
GTT GAA GCC AAG AAC GCT CTC GAG AAC TAC GCT TAC AAC ATG AGG AAC ACC ATC CAA GAC GAG AAG ATT GGT
GAG AAG CTC CCG GCT GCA GAC AAG AAG AAG ATC GAG GAT TCT ATT GAG CAG GCG ATT CAA TGG CTC GAG GGT
AAC CAG TTG GCT GAG GCT GAT GAG TTC GAA GAC AAG ATG AAG GAA TTG GAG AGC ATC TGC AAC CCA ATC ATT
GCC AAG ATG TAC CAA GGA GCT GGT GGT GAA GCC GGT GGT CCA GGT GCC TCT GGT ATG GAC GAT GAT GCT CCC
CCT GCT TCA GGC GGT GCT GGA CCT AAG ATC GAG GAG GTC GAC
**TAA**cggccgcactcgagatatctagacccagctttCTTGTACAAAGTGGTGCTAGGTGAG
TCTAGAGAGTTAATTAAGACCCGGGACTAGTCCCTAGAGTCCTGCTT.. ....GAGGCTCAG

***3. 35S::YFP:SVR (1813-1953)::OCS***

TGAGACTTT......ACAATTACC **ATG** GTG AGC AAG...... CTG TAC AAG TCC GGA CTC AGA TCT CGA GCT CAA GCT
TCG AAT TCT GCA GTC GAC GGT ACC GCG GGC CCG GGA TCA TCA ACA AGT Ttg tac aaa aaa gca ggc tcc acc
atg gga acc aat tca gtc gac tgg atc cgc ATG TCG GGT AAA GGA GAA GGA CCA GCT ATC GGT ATC GAT CTT
GGT ACC ACT TAC TCT TGC GTC GGA GTA TGG CAA CAC GAC CGT GTT GAG ATC ATT GCT AAT GAT CAA GGA AAC
AGA ACC ACG CCA TCT TAC GTT GCT TTC ACC GAC TCC GAG AGG TTG ATC GGT GAC GCA GCT AAG AAT CAG GTC
GCC ATG AAC CCC GTT AAC ACC GTT TTC GAC GCT AAG AGG TTG ATC GGT CGT CGT TTC TCT GAC AGC TCT GTT
CAG AGT GAC ATG AAA TTG TGG CCA TTC AAG ATT CAA GCC GGA CCT GCC GAT AAG CCA ATG ATC TAC GTC
GAA TAC AAG GGT GAA GAG AAA GAG TTC GCA GCT GAG GAG ATT TCT TCC ATG GTT CTT ATT AAG ATG CGT GAG
ATT GCT GAG GCT TAC CTT GGT GTC ACA ATC AAG AAC GCC GTT GTT ACC GTT CCA GCT TAC TTC AAC GAC TCT
CAG CGT CAG GCT ACA AAG GAT GCT GGT GTC ATC GCT GGT TTG AAC GTT ATG CGA ATC ATC AAC GAG CCT ACA
GCC GCC GCT ATT GCC TAC GGT CTT GAC AAA AAG GCT ACC AGC GTT GGA GAG AAG AAT GTT CTT ATC TTC GAT
CTT GGT GGT GGC ACT TTT GAT GTC TCT CTT CTT ACC ATT GAA GAG GGT ATC TTT GAG GTG AAG GCA ACT GCT
GGT GAC ACC CAT CTT GGT GGG GAA GAT TTT GAC AAC AGA ATG GTT AAC CAC TTT GTC CAA GAG TTC AAG AGG
AAG AGT AAG AAG GAT ATC ACC GGT AAC CCA AGA GCT CTT AGG AGG TTG AGA ACT TCC TGT GAG AGA GCG AAG
AGG ACT CTT TCT TCC ACT GCT CAG ACC ACC ATC GAG ATT GAC TCT CTA TAC GAG GGT ATC GAC TTC TAC TCC
ACC ATC ACC CGT GCT AGA TTT GAG GAG CTC AAC ATG GAT CTC TTC AGG AAG TGT ATG GAG CCA GTT GAG AAG
TGT CTT CGT GAT GCT AAG ATG GAC AAG AGC ACT GTT CAT GAT GTT GTC CTT GTT GGT GGT TCT ACC CGT ATC
CCT AAG GTT CAG CAA TTG CTC CAG GAC TTC TTC AAC GGC AAA GAG CTT TGC AAG TCT ATT AAC CCT GAT GAG
GCT GTT GCC TAC GGT GCT GCT GTC CAG GGA GCT ATT CTC AGC GGT GAA GGA AAC GAG AAG GTT CAA GAT CTT
CTA TTG CTC GAT GTC ACT CCT CTC TCC CTT GGT TTG GAA ACT GCC GGT GGT GTC ATG ACC ACT TTG ATC CCA
AGG AAC ACA ACC ATC CCA ACC AAG AAG GAA CAA GTC TTC TCC ACC TAC TCA GAC AAC CAA CCC GGT GTG TTG
ATC CAG GTG TAC GAA GGA GAG AGA GCC AGA ACC AAG GAC AAC AAC CTT CTT GGT AAA TTT GAG CTC TCC GGA
ATT CCT CCA GCT CCT CGT GGT GTC CCC CAG ATC ACA GTC TGC TTT GAC ATT GAT GCC AAT GGT ATC CTC AAT
GTC TCT GCT GAG GAC AAG ACC ACC GGA CAG AAG AAC AAG ATC ACC ATC ACC AAT GAC AAG GGT CGT CTC TCC
AAG GAT GAG ATT GAG AAG ATG GTT CAA GAG GCT GAG AAG TAC AAG TCC GAA GAC GAG GAG CAC AAG AAG AAG
GTT GAA GCC AAG AAC GCT CTC GAG AAC TAC GCT TAC AAC ATG AGG AAC ACC ATC CAA GAC GAG AAG ATT GGT
GAG AAG CTC CCG GCT GCA GAC AAG AAG AAG ATC GAG GAT TCT ATT GAG CAG GCG ATT CAA TGG CTC GAG GGT
AAC CAG TTG GCT GAG GCT GAT GAG TTC GAA GAC AAG ATG AAG GAA TTG GAG AGC ATC TGC AAC CCA ATC ATT
GCC AAG ATG TAC CAA GGA GCT GGT GGT GAA GCC GGT GGT CCA GGT GCC TCT GGT ATG GAC GAT GAT GCT CCC
CCT GCT TCA GGC GGT GCT GGA CCT AAG ATC GAG GAG GTC GAC **TAA**cggccgcactcgagatatctagacccagctttCTTGTA
CAAAGTGGTGCTAGGTGA GTCTAGAGAGTTAATTAAGACCCGGGACTAGTCCCTAGAGTCCTGCTT......GAGGCTCAG

***4. 35S::YFP:HSC70.1M (1-1953)::OCS (1-1191 original HSC70.1 sequence, 1192-1953 codon usage modified sequence)***

TGAGACTTT......ACAATTACC **ATG** GTG AGC AAG...... CTG TAC AAG TCC GGA CTC AGA TCT CGA GCT CAA GCT
TCG AAT TCT GCA GTC GAC GGT ACC GCG GGC CCG GGA TCA TCA ACA AGT Ttg tac aaa aaa gca ggc tcc acc
atg gga acc aat tca gtc gac tgg atc cgc ATG TCG GGT AAA GGA GAA GGA CCA GCT ATC GGT ATC GAT CTT
GGT ACC ACT TAC TCT TGC GTC GGA GTA TGG CAA CAC GAC CGT GTT GAG ATC ATT GCT AAT GAT CAA GGA AAC
AGA ACC ACG CCA TCT TAC GTT GCT TTC ACC GAC TCC GAG AGG TTG ATC GGT GAC GCA GCT AAG AAT CAG GTC
GCC ATG AAC CCC GTT AAC ACC GTT TTC GAC GCT AAG AGG TTG ATC GGT CGT CGT TTC TCT GAC AGC TCT GTT
CAG AGT GAC ATG AAA TTG TGG CCA TTC AAG ATT CAA GCC GGA CCT GCC GAT AAG CCA ATG ATC TAC GTC
GAA TAC AAG GGT GAA GAG AAA GAG TTC GCA GCT GAG GAG ATT TCT TCC ATG GTT CTT ATT AAG ATG CGT GAG
ATT GCT GAG GCT TAC CTT GGT GTC ACA ATC AAG AAC GCC GTT GTT ACC GTT CCA GCT TAC TTC AAC GAC TCT
CAG CGT CAG GCT ACA AAG GAT GCT GGT GTC ATC GCT GGT TTG AAC GTT ATG CGA ATC ATC AAC GAG CCT ACA
GCC GCC GCT ATT GCC TAC GGT CTT GAC AAA AAG GCT ACC AGC GTT GGA GAG AAG AAT GTT CTT ATC TTC GAT
CTT GGT GGT GGC ACT TTT GAT GTC TCT CTT CTT ACC ATT GAA GAG GGT ATC TTT GAG GTG AAG GCA ACT GCT
GGT GAC ACC CAT CTT GGT GGG GAA GAT TTT GAC AAC AGA ATG GTT AAC CAC TTT GTC CAA GAG TTC AAG AGG
AAG AGT AAG AAG GAT ATC ACC GGT AAC CCA AGA GCT CTT AGG AGG TTG AGA ACT TCC TGT GAG AGA GCG AAG
AGG ACT CTT TCT TCC ACT GCT CAG ACC ACC ATC GAG ATT GAC TCT CTA TAC GAG GGT ATC GAC TTC TAC TCC
ACC ATC ACC CGT GCT AGA TTT GAG GAG CTC AAC ATG GAT CTC TTC AGG AAG TGT ATG GAG CCA GTT GAG AAG
TGT CTT CGT GAT GCT AAG ATG GAC AAG AGC ACT GTT CAT GAT GTT GTC CTT GTT GGT GGT TCT ACC CGT ATC
CCT AAG GTT CAG CAA TTG CTC CAG GAC TTC TTC AAC GGC AAA GAG CTT TGC AAG TCT ATT AAC CCT GAT GAG

GCT GTT GCC TAC GGT GCT GCT GTC CAG GGA GCT ATT CTC AGC GGT GAA GGA AAC GAG AAG GTT CAA GAT CTT
CTT CTT CTT GAT GTT ACT CCT CTT TCT CTT GGA CTT GAA ACT GCT GGA GGA GTT ATG ACT ACT CTT ATT CCT
AGA AAT ACT ACT ATT CCT ACT AAG AAG GAA CAA GTT TTT TCT ACT TAT TCT GAT AAT CAA CCT GGA GTT CTT
ATT CAA GTT TAT GAA GGA GAA AGA GCT AGA ACT AAG GAT AAT AAT CTT CTT GGA AAG TTT GAA CTT TCT GGA
ATT CCT CCT GCT CCT AGA GGA GTT CCT CAA ATT ACT GTT TGT TTT GAT ATT GAT GCT AAT GGA ATT CTT AAT
GTT TCT GCT GAA GAT AAG ACT ACT GGA CAA AAG AAT AAG ATT ACT ATT ACT AAT GAT AAG GGA AGA CTT TCT
AAG GAT GAA ATT GAA AAG ATG GTT CAA GAA GCT GAA AAG TAT AAG TCT GAA GAT GAA GAA CAT AAG AAG AAG
GTT GAA GCT AAG AAT GCT CTT GAA AAT TAT GCT TAT AAT ATG AGA AAT ACT ATT CAA GAT GAA AAG ATT GGA
GAA AAG CTT CCT GCT GCT GAT AAG AAG AAG ATT GAA GAT TCT ATT GAA CAA GCT ATT CAA TGG CTT GAA GGA
AAT CAA CTT GCT GAA GCT GAT GAA TTT GAA GAT AAG ATG AAG GAA CTT GAA TCT ATT TGT AAT CCT ATT ATT
GCT AAG ATG TAT CAA GGA GCT GGA GGA GAA GCT GGA GGA CCT GGA GCT TCT GGA ATG GAT GAT GAT GCT CCT
CCT GCT TCT GGA GGA GCT GGA CCT AAG ATT GAA GAA GTT GAT
**TAA**cggccgcactcgagatatctagacccagctttCTTGTACAAAGTGGTGCTAGGTGAGTCTAGAGAGTT
AATTAAGACCCGGGACTAGTCCCTAGAGTCCTGCTT......GAGGCTCAG

***5. 35S::YFP:HSC70.1M ΔSVR (1-1812)::OCS (1-1191 original HSC70.1 sequence, 1192-1953 codon usage modified sequence)***

TGAGACTTT......ACAATTACC **ATG** GTG AGC AAG...... CTG TAC AAG TCC GGA CTC AGA TCT CGA GCT CAA GCT
TCG AAT TCT GCA GTC GAC GGT ACC GCG GGC CCG GGA TCA TCA ACA AGT Ttg tac aaa aaa gca ggc tcc acc
atg gga acc aat tca gtc gac tgg atc cgc ATG TCG GGT AAA GGA GAA GGA CCA GCT ATC GGT ATC GAT CTT
GGT ACC ACT TAC TCT TGC GTC GGA GTA TGG CAA CAC GAC CGT GTT GAG ATC ATT GCT AAT GAT CAA GGA AAC
AGA ACC ACG CCA TCT TAC GTT GCT TTC ACC GAC TCC GAG AGG TTG ATC GGT GAC GCA GCT AAG AAT CAG GTC
GCC ATG AAC CCC GTT AAC ACC GTT TTC GAC GCT AAG AGG TTG ATC GGT CGT CGT TTC TCT GAC AGC TCT GTT
CAG AGT GAC ATG AAA TTG TGG CCA TTC AAG ATT CAA GCC GGA CCT GCC GAT AAG CCA ATG ATC TAC GTC
GAA TAC AAG GGT GAA GAG AAA GAG TTC GCA GCT GAG GAG ATT TCT TCC ATG GTT CTT ATT AAG ATG CGT GAG
ATT GCT GAG GCT TAC CTT GGT GTC ACA ATC AAG AAC GCC GTT GTT ACC GTT CCA GCT TAC TTC AAC GAC TCT
CAG CGT CAG GCT ACA AAG GAT GCT GGT GTC ATC GCT GGT TTG AAC GTT ATG CGA ATC ATC AAC GAG CCT ACA
GCC GCC GCT ATT GCC TAC GGT CTT GAC AAA AAG GCT ACC AGC GTT GGA GAG AAG AAT GTT CTT ATC TTC GAT
CTT GGT GGT GGC ACT TTT GAT GTC TCT CTT CTT ACC ATT GAA GAG GGT ATC TTT GAG GTG AAG GCA ACT GCT
GGT GAC ACC CAT CTT GGT GGG GAA GAT TTT GAC AAC AGA ATG GTT AAC CAC TTT GTC CAA GAG TTC AAG AGG
AAG AGT AAG AAG GAT ATC ACC GGT AAC CCA AGA GCT CTT AGG AGG TTG AGA ACT TCC TGT GAG AGA GCG AAG
AGG ACT CTT TCT TCC ACT GCT CAG ACC ACC ATC GAG ATT GAC TCT CTA TAC GAG GGT ATC GAC TTC TAC TCC
ACC ATC ACC CGT GCT AGA TTT GAG GAG CTC AAC ATG GAT CTC TTC AGG AAG TGT ATG GAG CCA GTT GAG AAG
TGT CTT CGT GAT GCT AAG ATG GAC AAG AGC ACT GTT CAT GAT GTT GTC CTT GTT GGT GGT TCT ACC CGT ATC
CCT AAG GTT CAG CAA TTG CTC CAG GAC TTC TTC AAC GGC AAA GAG CTT TGC AAG TCT ATT AAC CCT GAT GAG
GCT GTT GCC TAC GGT GCT GCT GTC CAG GGA GCT ATT CTC AGC GGT GAA GGA AAC GAG AAG GTT CAA GAT CTT
CTT CTT CTT GAT GTT ACT CCT CTT TCT CTT GGA CTT GAA ACT GCT GGA GGA GTT ATG ACT ACT CTT ATT CCT
AGA AAT ACT ACT ATT CCT ACT AAG AAG GAA CAA GTT TTT TCT ACT TAT TCT GAT AAT CAA CCT GGA GTT CTT
ATT CAA GTT TAT GAA GGA GAA AGA GCT AGA ACT AAG GAT AAT AAT CTT CTT GGA AAG TTT GAA CTT TCT GGA
ATT CCT CCT GCT CCT AGA GGA GTT CCT CAA ATT ACT GTT TGT TTT GAT ATT GAT GCT AAT GGA ATT CTT AAT
GTT TCT GCT GAA GAT AAG ACT ACT GGA CAA AAG AAT AAG ATT ACT ATT ACT AAT GAT AAG GGA AGA CTT TCT
AAG GAT GAA ATT GAA AAG ATG GTT CAA GAA GCT GAA AAG TAT AAG TCT GAA GAT GAA GAA CAT AAG AAG AAG
GTT GAA GCT AAG AAT GCT CTT GAA AAT TAT GCT TAT AAT ATG AGA AAT ACT ATT CAA GAT GAA AAG ATT GGA
GAA AAG CTT CCT GCT GCT GAT AAG AAG AAG ATT GAA GAT TCT ATT GAA CAA GCT ATT CAA TGG CTT GAA GGA
AAT CAA CTT GCT GAA GCT GAT GAA TTT GAA GAT AAG ATG AAG GAA CTT GAA TCT ATT TGT AAT CCT ATT ATT
GCT AAG ATG TAT CAA GGA GCT GGA GGA GAA GCT GGA GGA CCT GGA GCT TCT GGA ATG GAT GAT GAT GCT CCT
CCT GCT TCT GGA GGA GCT GGA CCT AAG ATT GAA GAA GTT GAT
**TAA**cggccgcactcgagatatctagacccagctttCTTGTACAAAGTGGTGCTAGGTGAGTCTAGAGAGTT
AATTAAGACCCGGGACTAGTCCCTAGAGTCCTGCTT......GAGGCTCAG

***6. 35S::YFP(s)::HSC70.1***

TGAGACTTT......ACAATTACC **ATG** GTG AGC AAG...... CTG TAC AAG TCC GGA CTC AGA TCT CGA GCT CAA GCT
TCG AAT TCT GCA GTC GAC GGT ACC GCG GGC CCG GGA TCA TCA ACA AGT Ttg tac aaa aaa gca ggc tcc acc
atg gga acc aat tca gtc gac tgg atc cgc **TAA** ATG TCG GGT AAA GGA GAA GGA CCA GCT ATC GGT ATC GAT
CTT GGT ACC ACT TAC TCT TGC GTC GGA GTA TGG CAA CAC GAC CGT GTT GAG ATC ATT GCT AAT GAT CAA GGA
AAC AGA ACC ACG CCA TCT TAC GTT GCT TTC ACC GAC TCC GAG AGG TTG ATC GGT GAC GCA GCT AAG AAT CAG
GTC GCC ATG AAC CCC GTT AAC ACC GTT TTC GAC GCT AAG AGG TTG ATC GGT CGT CGT TTC TCT GAC AGC TCT
GTT CAG AGT GAC ATG AAA TTG TGG CCA TTC AAG ATT CAA GCC GGA CCT GCC GAT AAG CCA ATG ATC TAC GTC
GAA TAC AAG GGT GAA GAG AAA GAG TTC GCA GCT GAG GAG ATT TCT TCC ATG GTT CTT ATT AAG ATG CGT GAG
ATT GCT GAG GCT TAC CTT GGT GTC ACA ATC AAG AAC GCC GTT GTT ACC GTT CCA GCT TAC TTC AAC GAC TCT
CAG CGT CAG GCT ACA AAG GAT GCT GGT GTC ATC GCT GGT TTG AAC GTT ATG CGA ATC ATC AAC GAG CCT ACA
GCC GCC GCT ATT GCC TAC GGT CTT GAC AAA AAG GCT ACC AGC GTT GGA GAG AAG AAT GTT CTT ATC TTC GAT
CTT GGT GGT GGC ACT TTT GAT GTC TCT CTT CTT ACC ATT GAA GAG GGT ATC TTT GAG GTG AAG GCA ACT GCT
GGT GAC ACC CAT CTT GGT GGG GAA GAT TTT GAC AAC AGA ATG GTT AAC CAC TTT GTC CAA GAG TTC AAG AGG
AAG AGT AAG AAG GAT ATC ACC GGT AAC CCA AGA GCT CTT AGG AGG TTG AGA ACT TCC TGT GAG AGA GCG AAG
AGG ACT CTT TCT TCC ACT GCT CAG ACC ACC ATC GAG ATT GAC TCT CTA TAC GAG GGT ATC GAC TTC TAC TCC
ACC ATC ACC CGT GCT AGA TTT GAG GAG CTC AAC ATG GAT CTC TTC AGG AAG TGT ATG GAG CCA GTT GAG AAG
TGT CTT CGT GAT GCT AAG ATG GAC AAG AGC ACT GTT CAT GAT GTT GTC CTT GTT GGT GGT TCT ACC CGT ATC
CCT AAG GTT CAG CAA TTG CTC CAG GAC TTC TTC AAC GGC AAA GAG CTT TGC AAG TCT ATT AAC CCT GAT GAG
GCT GTT GCC TAC GGT GCT GCT GTC CAG GGA GCT ATT CTC AGC GGT GAA GGA AAC GAG AAG GTT CAA GAT CTT
CTA TTG CTC GAT GTC ACT CCT CTC TCC CTT GGT TTG GAA ACT GCC GGT GGT GTC ATG ACC ACT TTG ATC CCA
AGG AAC ACA ACC ATC CCA ACC AAG AAG GAA CAA GTC TTC TCC ACC TAC TCA GAC AAC CAA CCC GGT GTG TTG
ATC CAG GTG TAC GAA GGA GAG AGA GCC AGA ACC AAG GAC AAC AAC CTT CTT GGT AAA TTT GAG CTC TCC GGA
ATT CCT CCA GCT CCT CGT GGT GTC CCC CAG ATC ACA GTC TGC TTT GAC ATT GAT GCC AAT GGT ATC CTC AAT
GTC TCT GCT GAG GAC AAG ACC ACC GGA CAG AAG AAC AAG ATC ACC ATC ACC AAT GAC AAG GGT CGT CTC TCC
AAG GAT GAG ATT GAG AAG ATG GTT CAA GAG GCT GAG AAG TAC AAG TCC GAA GAC GAG GAG CAC AAG AAG AAG

GTT GAA GCC AAG AAC GCT CTC GAG AAC TAC GCT TAC AAC ATG AGG AAC ACC ATC CAA GAC GAG AAG ATT GGT GAG AAG CTC CCG GCT GCA GAC AAG AAG AAG ATC GAG GAT TCT ATT GAG CAG GCG ATT CAA TGG CTC GAG GGT AAC CAG TTG GCT GAG GCT GAT GAG TTC GAA GAC AAG ATG AAG GAA TTG GAG AGC ATC TGC AAC CCA ATC ATT GCC AAG ATG TAC CAA GGA GCT GGT GGT GAA GCC GGT GGT CCA GGT GCC TCT GGT ATG GAC GAT GAT GCT CCC CCT GCT TCA GGC GGT GCT GGA CCT AAG ATC GAG GAG GTC GAC

**TAA**cggccgcactcgagatatctagacccagctttCTTGTACAAAGTGGTGCTAG

GTGAGTCTAGAGAGTTAATTAAGACCCGGGACTAGTCCCTAGAGTCCTGC TT......GAGGCTCAG

#### *6. Sequence alignment of wild-type HSC70.1 vs. modified HSC70.1 CDS (HSC70.1M)*

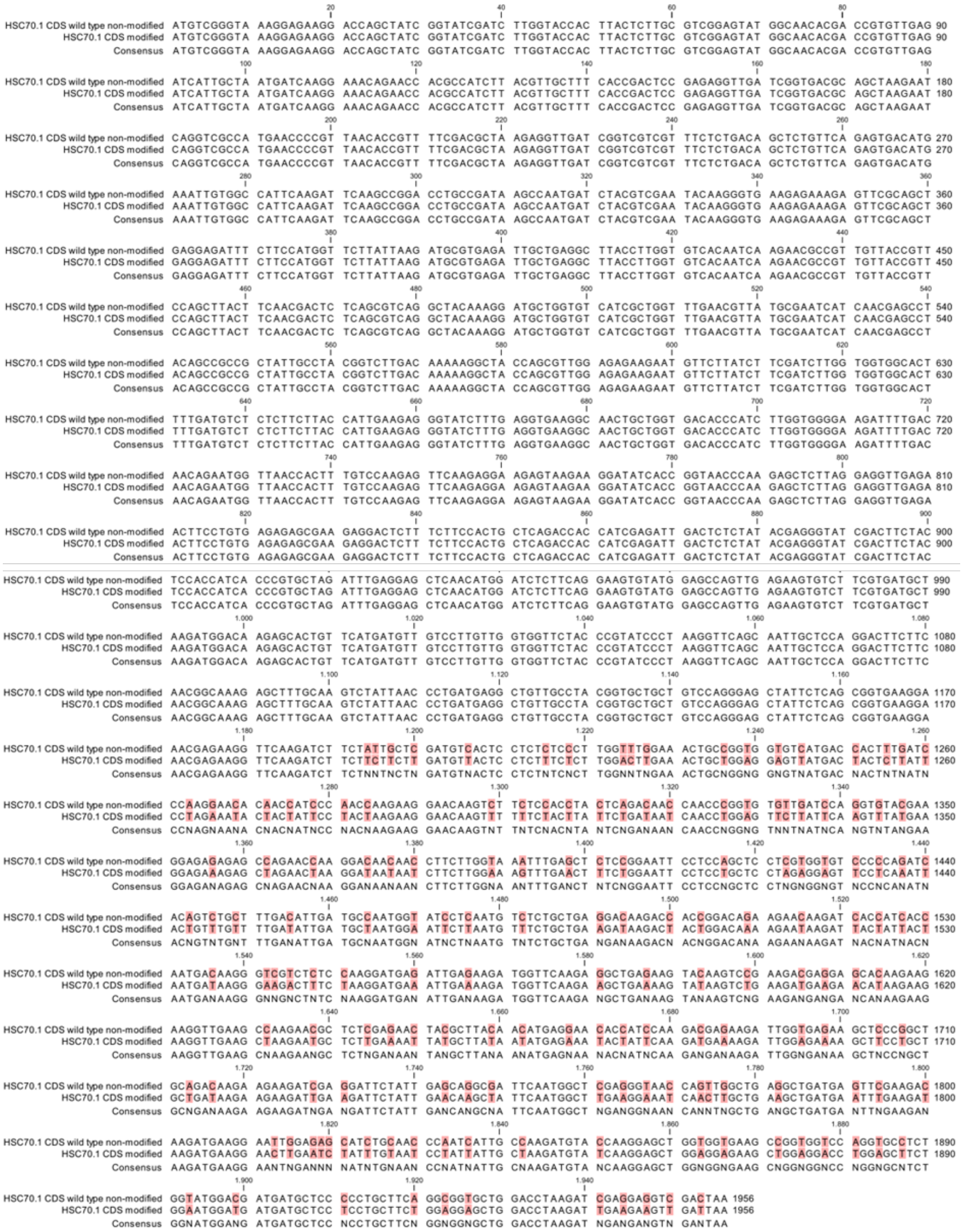

#### *7.* Sequence alignment of wild-type HSC70.1 vs. modified HSC70.1 (HSC70.1M) aa sequences

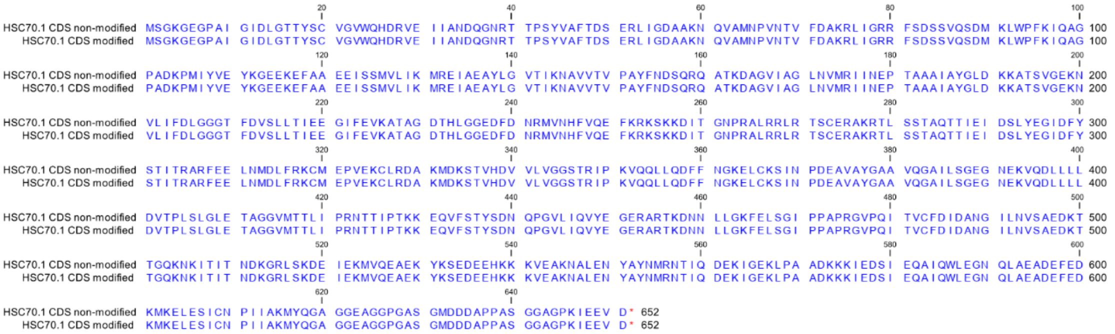

### Supplementary Dataset S2

#### RT and PCR Primers used in this study.

| Purpose / name of primer | Primer sequence | Size of PCR fragment |
|---|---|---|
| Construction *of YFP-HSC70.1 and YFP-HSC70.1 ΔSVR* fusion forward primer | FK1527-F 5'-GCGGGATCCGCATGTCGGGTAAAGGAGAAG-3' | |
| Construction of *YFP-HSC70.1* and *YFP-HSC70.1 SVR* fusion reverse primer | FK1528-R 5'-GCGCGGCCGTTAGTCGACCTCCTCGATC-3' | Size of *YFP-HSC70.1* full length 1956 bp |
| Construction *of YFP-HSC70.1 ΔSVR'* fusion *r*everse primer | FK1529-R 5'-GCGCGGCCGTTATTCCTTCATCTTGTCTTCG-3' | Size of *YFP-TCTP1 ΔSVR'* 1815 bp |
| Construction *of YFP-HSC70.1 SVR fusion* forward primer | FK1530-F 5'-GCGCGGCCGTTAGTCGACCTCCTCGATC- 3' | Size of *YFP-HSC70.1 SVR* 143 bp |
| *YFP-HSC70.1, YFP-HSC70.1 ΔSVR, YFP-HSC70.1M* and *YFP-HSC70.1M ΔSVR* RT detection | FK938-F 5'-CCCGACAACCACTACCTGAG- 3' | 501 bp |
| | FK1531-R 5'-GAAACGACGACCGATCAAC- 3' | |
| *YFP-HSC70.1 SVR* RT detection primer | FK938-F 5'-CCCGACAACCACTACCTGAG- 3' | 405bp |
| | FK1532-R 5'-TTAGTCGACCTCCTCGATCTTAGG-3' | |
| *HSC70.1* RT detection | FK1533-F 5'-CCGATAAGCCAATGATCTACGTC-3' | |
| | FK1534-R 5'-GGCGGCTGTAGGCTCGTTG-3' | 245bp |
| *ACTIN2* (At3g18780) detection | FK424-F 5'-GGAAGGATCTGTACGGTAAC-3' | |
| | FK425-R 5'-TGTGAACGATTCCTGGACCT-3' | 245 bp |
| *HSC70.1* qRT-PCR detection | FK1330-F 5'-GCTGGTGGTGAAGCCGGTG-3' | |
| | FK1331-R 5'-GGAGAAAGAGAGAGGTCAATGTC-3' | 129 bp |
| *HSC70.4* qRT detection primer | FK1214-F 5'-CTACATGTCCCACTTGCCTG-3' | |
| | FK1215-R 5'-CTCGGGGTAGTTGCCAGAT -3' | 277 bp |
| *UBQ10* qRT detection primer | FK1095-F 5'-CACACTTCACTTGGTCTTGCGT-3 | |
| | FK1096-R 5'-TAGTCTTTCCGGTGAGAGTCTTCA-3' | 70 bp |
| *hsc70.1* SALK_135531 T-DNA Insertion primer | FK1748-F 5'-AAGGAGAAGGACCAGCTATCG-3 | |
| | FK1749-R 5'-TCTTCGCTCTCTCACAGGAAG-3' | 1146 bp |
| *hsc70.4* SALK_088253 T-DNA Insertion primer | FK1750-F 5'-CCAAATACGAAGCCACTTGAG-3 | |
| | FK11751-R 5'-TACCGAAGACGGTGTTGGTAG-3' | 1133 bp |
| *HSC70.1* RIP primer 1 | FK1520-F 5'-CTTACAACATGAGGAACACCATCC-3' | |
| | FK1521-R 5'-CATCGTCCATACCAGAGGCAC-3' | 249 bp |
| *HSC70.*1 RIP primer 2 | FK1533-F 5'-AATGACAAGGGTCGTCTCTCC-3' | |
| | FK1534-R 5'-CTCGAGCCATTGAATCGCC-3' | 234 bp |
| *HSC70.*1 RIP primer 3 | FK1752-F 5'-GGAGAAAAGCTTCCTGCTGCTG-3' | |

| | | |
|---|---|---|
| | FK1753-R 5’-TAGGTCCAGCTCCTCCAGAAGC-3’ | 244 bp |
| *BAG1* RIP primer 1 | FK1657-F 5’- GGACCATTTGTGTTAGATTCTTCTGC -3’<br><br>FK1658-R 5’-GCGGGATCGCGTTATTTTTC-3’ | 101bp |
| *T7 YFP* primer | 5’-TAATACGACTCACTATAGATGGTGAGCAAGGGCGAGGAGC TGTTCACC-3’<br><br>5’-TTATTAGTACAGCTCGTCCATGCCGAG-3’ | 717 bp |
| *T7 YFP-HSC70.1* primer | 5’-TAATACGACTCACTATAGATGGTGAGCAAGGGCGAGGAGC TGTTCACC-3’<br><br>5’-TTA TTA GTCGACCTCCTCGATCTTAGG-3’ | 2820 bp |
