## Supplemental Tables and Figures for "Non-cell-autonomous HSC70.1 chaperone displays homeostatic feed-back regulation by binding its own mRNA"

### Supplementary Figure S1
### Yang et al. 2022

**Predicted RNA folding structures of used *HSC70.1 CDS* sequences**

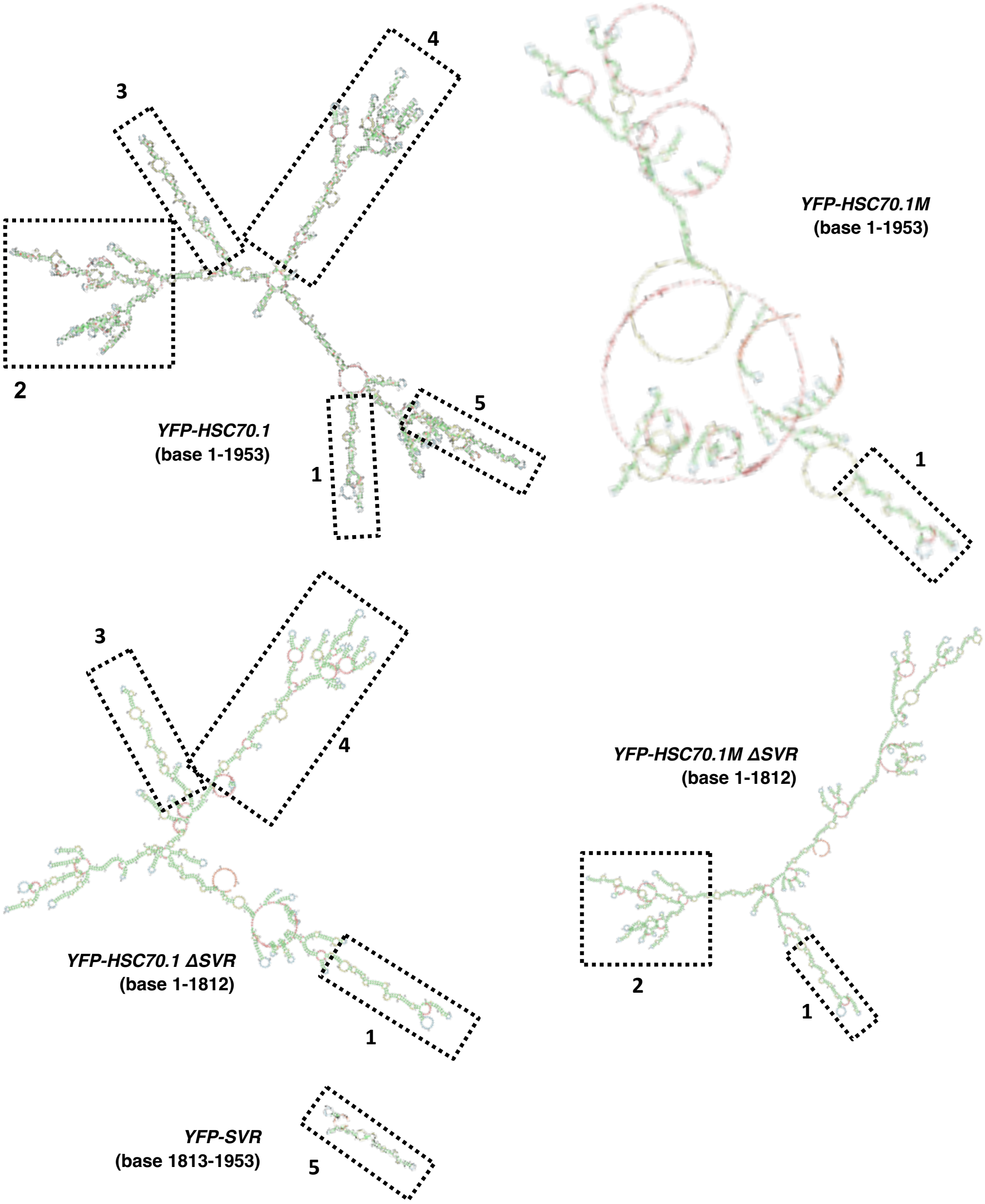


**Figure S1. Schematic predicted folding structures of used *HSC70.1* coding sequences (CDS) predicted according to their Minimal Free Energy (MFE) (Zuker et al., 1981; Lorenz et al., 2011)** Black boxes indicate five conserved structures. Note that deletion of the relative stable SVR region does not seem to affect the 5' folding of *HSC70.1* region 1, 3 and 4, and seems to slightly change region 2.

### Supplementary Figure S2
### Yang et al. 2022

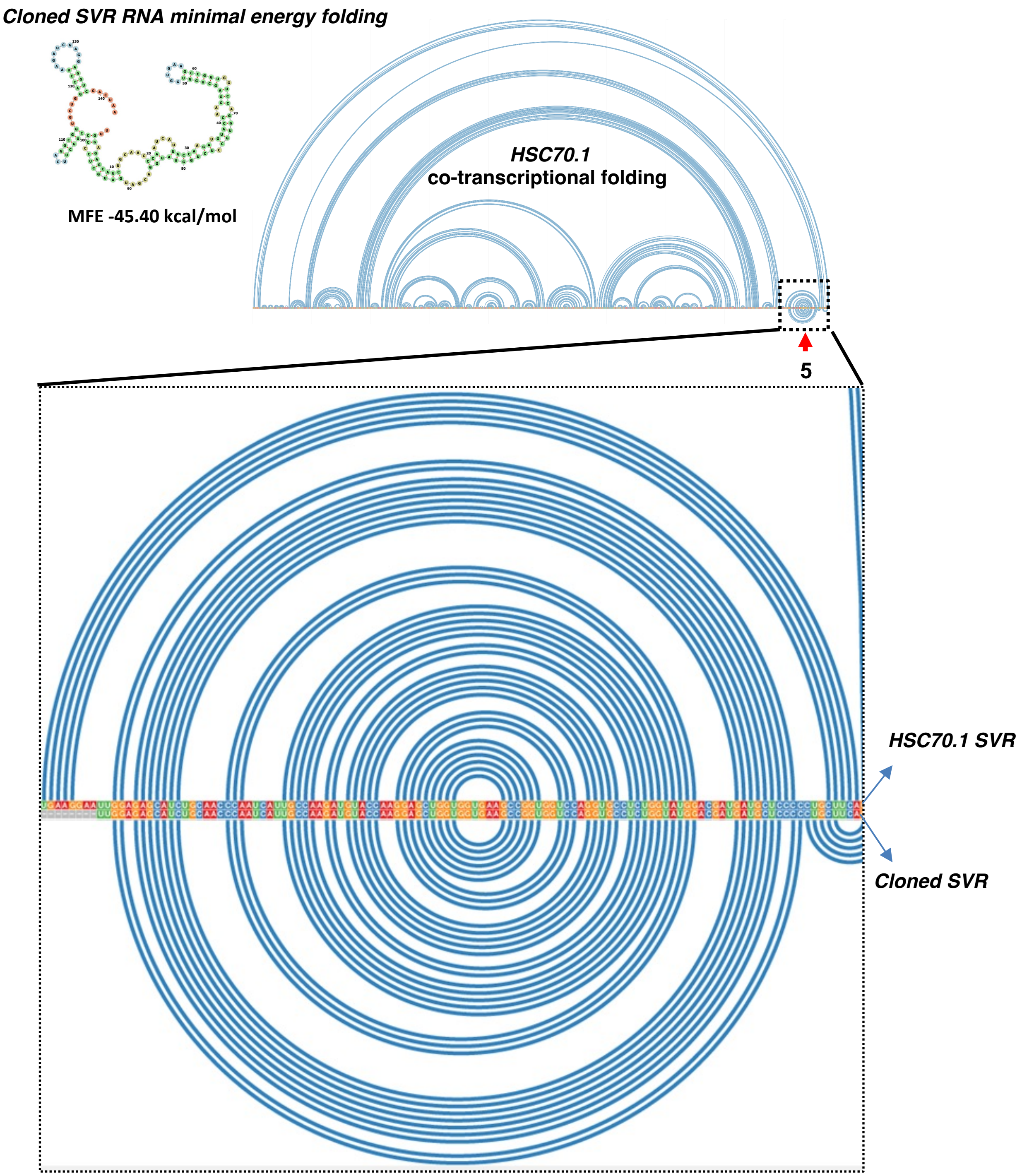


**Figure S2. Comparison of the hydrogen bonds connection between *HSC70.1* and cloned *SVR* CDS nucleotides predicted according to their co-transcriptional folding (Co-Fold) (Proctor et al. 2012).** Note that the hydrogen bonds connections of SVR region is highly stable.

### Supplementary Figure S3
### Yang et al. 2022

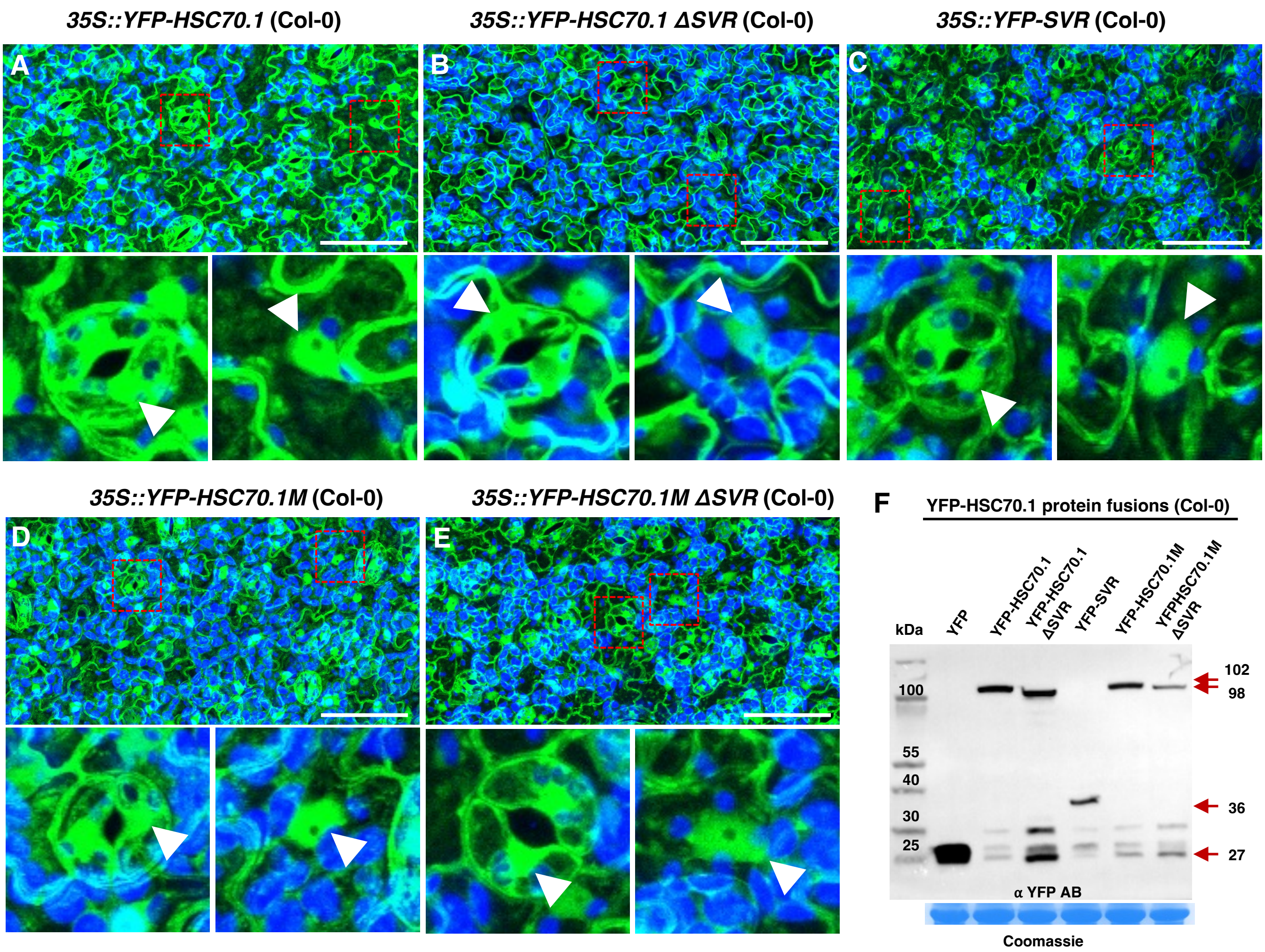


**Figure S3. Confocal laser scanning microscopy (CLSM) images of epidermal cells expressing the fusion constructs in Arabidopsis.**

A to E, All YFP fusions expressing a full-length *HSC70.1* or truncated *HSC70.1* or modified sequence (*HSC70.1M* and *HSC70.1MΔSVR*) were detected in both the cytoplasm and the nuclei. Blue color: chloroplastic auto-fluorescence. Green color: YFP fluorescence. Bar: 50 $\mu$m.

F, Western-blot detection of YFP fusion proteins produced in two-week-old transgenic Arabidopsis (Col-0) seedlings by GFP/YFP specific antibody. Calculated protein sizes including linker sequences are: YFP 27 kDa, YFP-HSC70.1 102 kDa, YFP-HSC70.1ΔSVR 98 kDa, YFP-SVR 36 kDa, YFP-HSC70.1M 102 kDa, YFP-HSC70.1MΔSVR 98 kDa. Red arrows mark apparent sizes of the YFP fusions. Lower panel: loading control.

### Supplementary Figure S4
### Yang et al. 2022

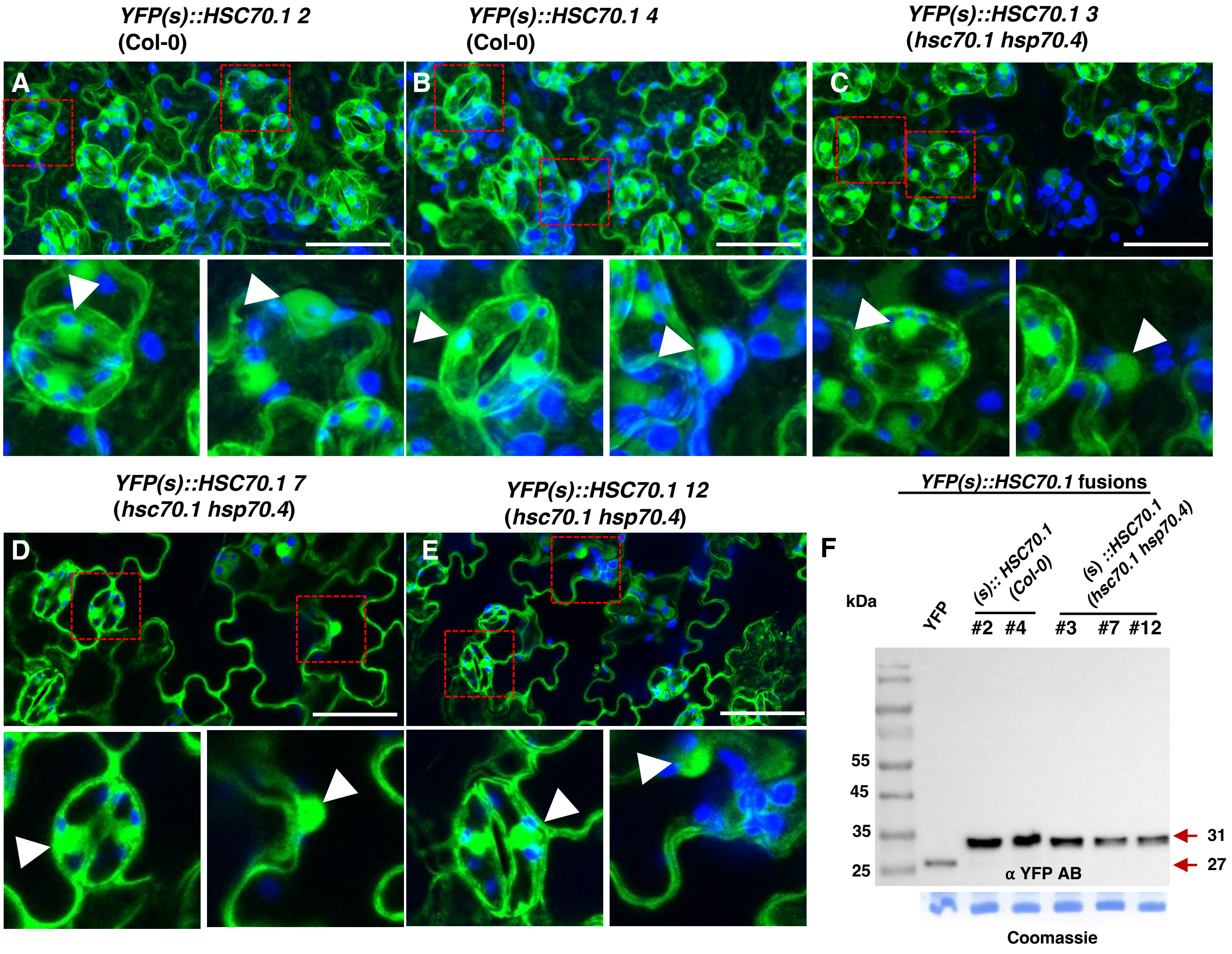


**Figure S4. Localization and expression of Y*FP(s)::HSC70.1* fusion construct in Col-0 wild-type and *hsc70.1 hsp70.4* mutants.**

A and B, Confocal laser scanning microscopy (CLSM) images of transgenic *YFP(s)::HSC70.1* Col-0 lines.

C to E, CLSM images of transgenic *YFP(s)::HSC70.1 hsc70.1 hsp70.4* lines. In both wild type and mutant lines the YFP protein was detected in cytoplasm and nuclei. Blue color: plastid auto-fluorescence. Green color: YFP fluorescence. Bar: 10 $\mu$m.

F, Western-blot detection of YFP(s)::HSC70.1 fusion protein produced in two-week-old transgenic Col-0 and *hsc70.1 hsp70.4* mutants. Lower panel: loading control. Red arrows indicated expected size of approx. 31 kDa due to the presence of linker sequences before the introduced stop codon.

### Supplementary Figure S5
### Yang et al. 2022

#### Identified transcripts enriched in YFP-HSC70.1 RIP assays

| TAIR # | Fold change | $Log_2$ fold change | P-value | FDR p-value | Name | RNA binding activity of encoded protein | Graft mobile (Col/Ped dtataset mobility blue) |
|---|---|---|---|---|---|---|---|
| AT1G21310.1 | 4.10 | 2.03 | 7.02E-4 | 0.01 | AT1G21310.1 \| extensin 3 | na | na |
| AT1G56070.1 | 3.87 | 1.95 | 2.33E-3 | 0.03 | AT1G56070.1 \| Ribosomal protein S5, LOS1 | yes | na |
| AT1G59870.1 | 7.24 | 2.86 | 1.38E-3 | 0.02 | AT1G59870.1 \| PEN3 | yes | yes |
| AT1G77760.1 | 5.05 | 2.34 | 1.63E-3 | 0.02 | AT1G77760.1 \| nitrate reductase 1, NIA1 | na | na |
| AT2G18020.1 | 5.79 | 2.53 | 3.09E-3 | 0.04 | AT2G18020.1 \| Ribosomal protein L2 family | yes | na |
| AT2G20410.1 | 11.13 | 3.48 | 1.25E-4 | 3.72E-3 | RNA (U2.1), snRNA / UTR of AT2G20410.1 | yes | na |
| AT3G08580.1 | 4.93 | 2.30 | 3.96E-3 | 0.04 | AT3G08580.1 \| ADP/ATP carrier 1, AAC | yes | na |
| AT4G14040.1 | 10.01 | 3.32 | 8.26E-4 | 0.01 | AT4G14040.1 \| selenium-binding protein 2 | na | na |
| AT4G35090.1 | 3.17 | 1.67 | 1.35E-3 | 0.02 | AT4G35090.1 \| catalase 2 | yes | na |
| AT5G02500.1 | 3.36 | 1.75 | 4.94E-3 | 0.05 | AT5G02500.1 \| HSC70-1 | yes | yes |
| AT5G20010.1 | 5.10 | 2.35 | 4.45E-3 | 0.04 | AT5G20010.1 \| RAS-related nuclear protein-1 | yes | yes |
| AT5G35630.3 | 5.33 | 2.41 | 8.23E-4 | 0.01 | AT5G35630.3 \| glutamine synthetase 2 | yes | yes |
| AT5G52060.1 | 51.08 | 5.67 | 1.19E-5 | 6.10E-4 | AT5G52060.1 \| HSC70 cochaperone BCL-2-associated athanogene 1 | na | na |

Note that 9 of 13 enriched transcripts encode RNA binding proteins and that 4 of the 13 transcripts were annotated as mobile according to Thieme et al. 2015.

**Figure S5. Identified transcripts found enriched in the RIP assays.**

Three biological replicates of erYFP and YFP-HSC70 RIP samples were submitted to RNAseq and analysed by CLC genomics workbench 21 software against released Araport 11 CDS sequences using standard settings. 13 transcripts were found significantly enriched with an FDR p value ≤ 0.05.

### Supplementary Figure S6
### Yang et al. 2022

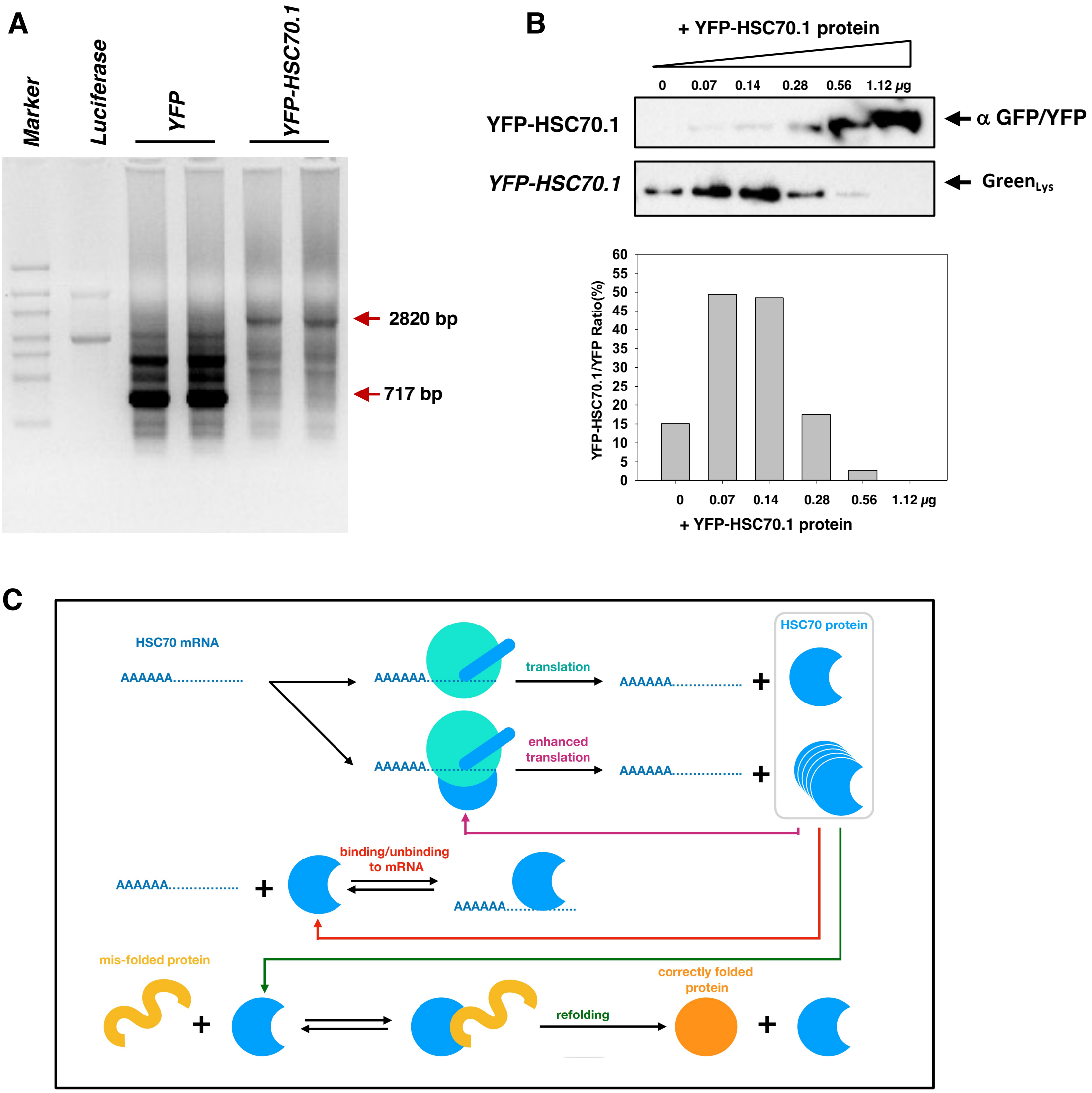


**Figure S6.** ***HSC70.1 in vitro*** **translation assay and reaction pathway used to calculate negative feedback of translation.**

A, Transcribed *YFP* and *YFP-HSC70.1* RNAs were analyzed by 1% agarose gel electrophoresis. Red arrows indicate *YFP-HSC70.1* and *YFP* RNA produced *in vitro* by T7 transcriptase.

B, Upper panel: Translation of YFP-HSC70.1 and YFP protein in the Wheat Germ Expression (WGE) system in the presence of increasing amounts of YFP-HSC70.1 extracted from 10 days old YFP-HSC70.1 transgenic plants. α -GFP/YFP AB and fluorescent Green$_{lys}$ was used to detected added YFP-HSC70.1 and its affect on *in vitro* translation, respectively. Lower Panel: Detected relative expression of YFP-HSC70.1 vs. YFP protein.

C, Reaction pathway used in calculating the model shown in Figure 2. The presence of HSC70 protein is assumed to enhance translation until a concentration is reached at which it binds to *HSC70* mRNA which blocks translation. HSC70 protein is assumed to bind more tightly to mis-folded protein than to its own mRNA and to assist in refolding of the client protein.

### Supplementary Figure S7
### Yang et al. 2022

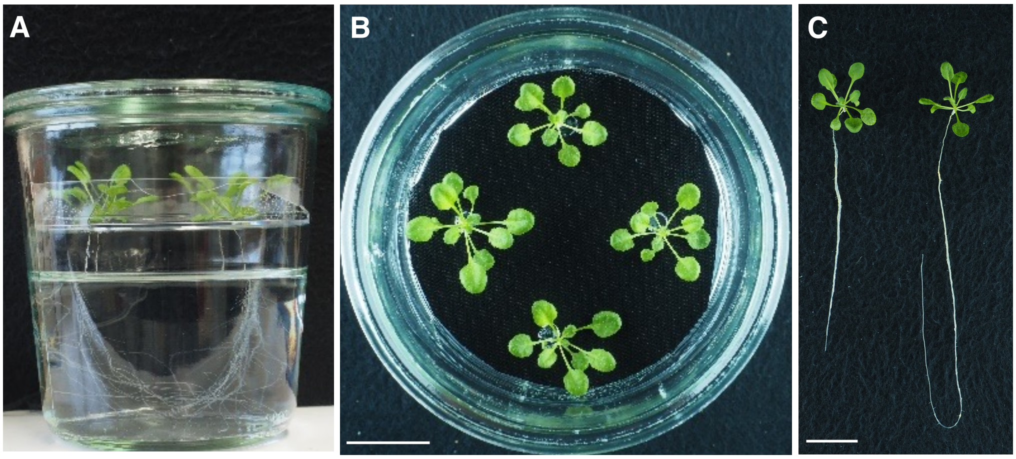


**Figure S7. Representative images of *YFP-HSC70.1* / Col-0 grafted plants grown in liquid culture 30 days after grafting.**

A and B Pictures of grafted plants grown in the hydroponic system.

C Picture of two grafted plants exemplifying the variation of growth detected in the liquid culture system. Bar: 2 cm.

### Supplementary Figure S8
### Yang et al. 2022

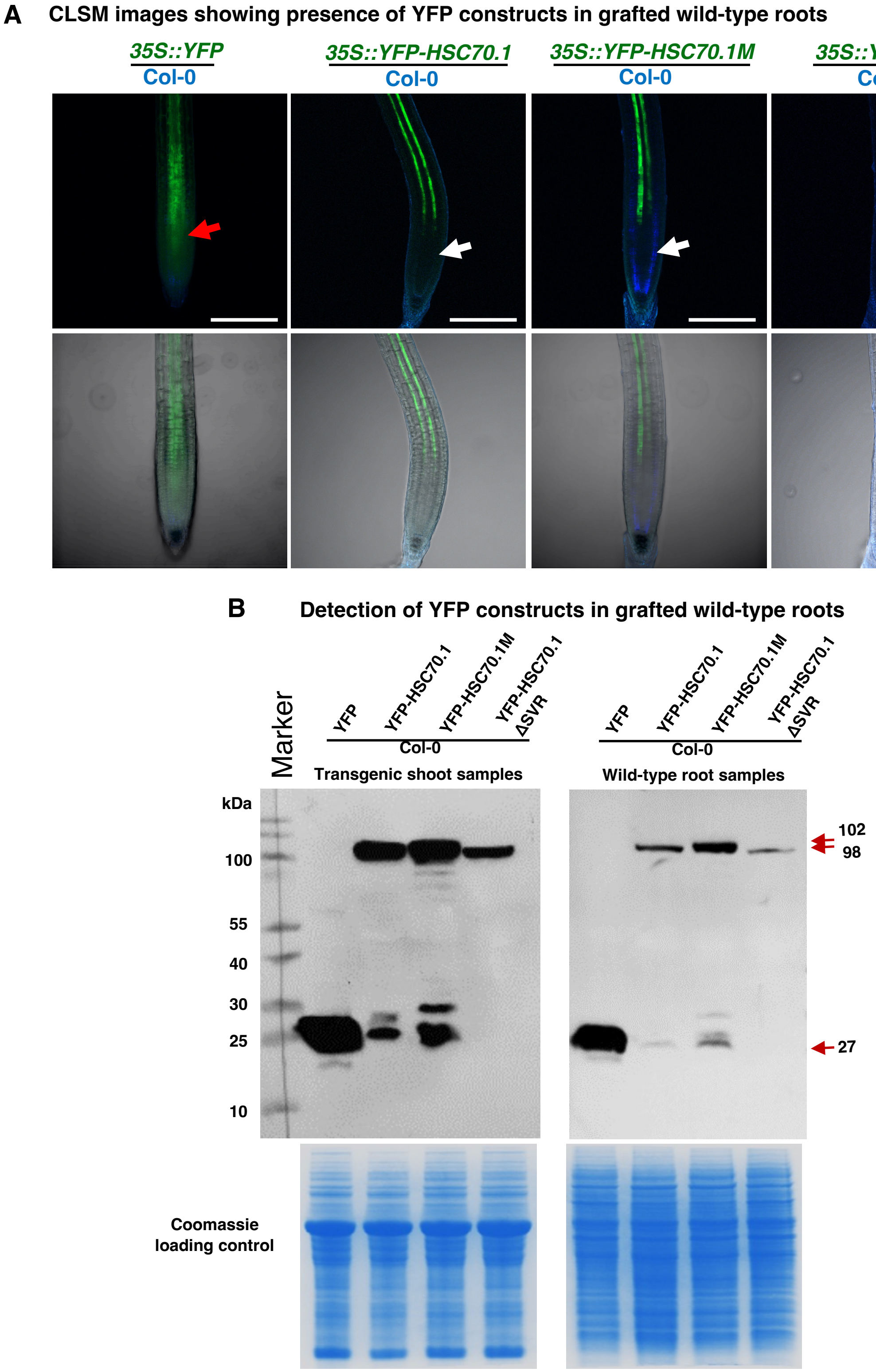


**Figure S8. Presence and integrity of YFP-fusion constructs in grafted wild-type roots.**

A, CLSM images of grafted wild-type roots showing YFP fluorescence in the phloem vasculature. Blue color: plastid auto-fluorescence. Green color: YFP fluorescence. Bar: 200 $\mu$m.

B, Western-blot assays confirming expression of full-length YFP fusion proteins in transgenic scions (left panel) and their presence in grafted wild-type (Col-0) roots (right panel).

### Supplementary Figure S9
### Yang et al. 2022

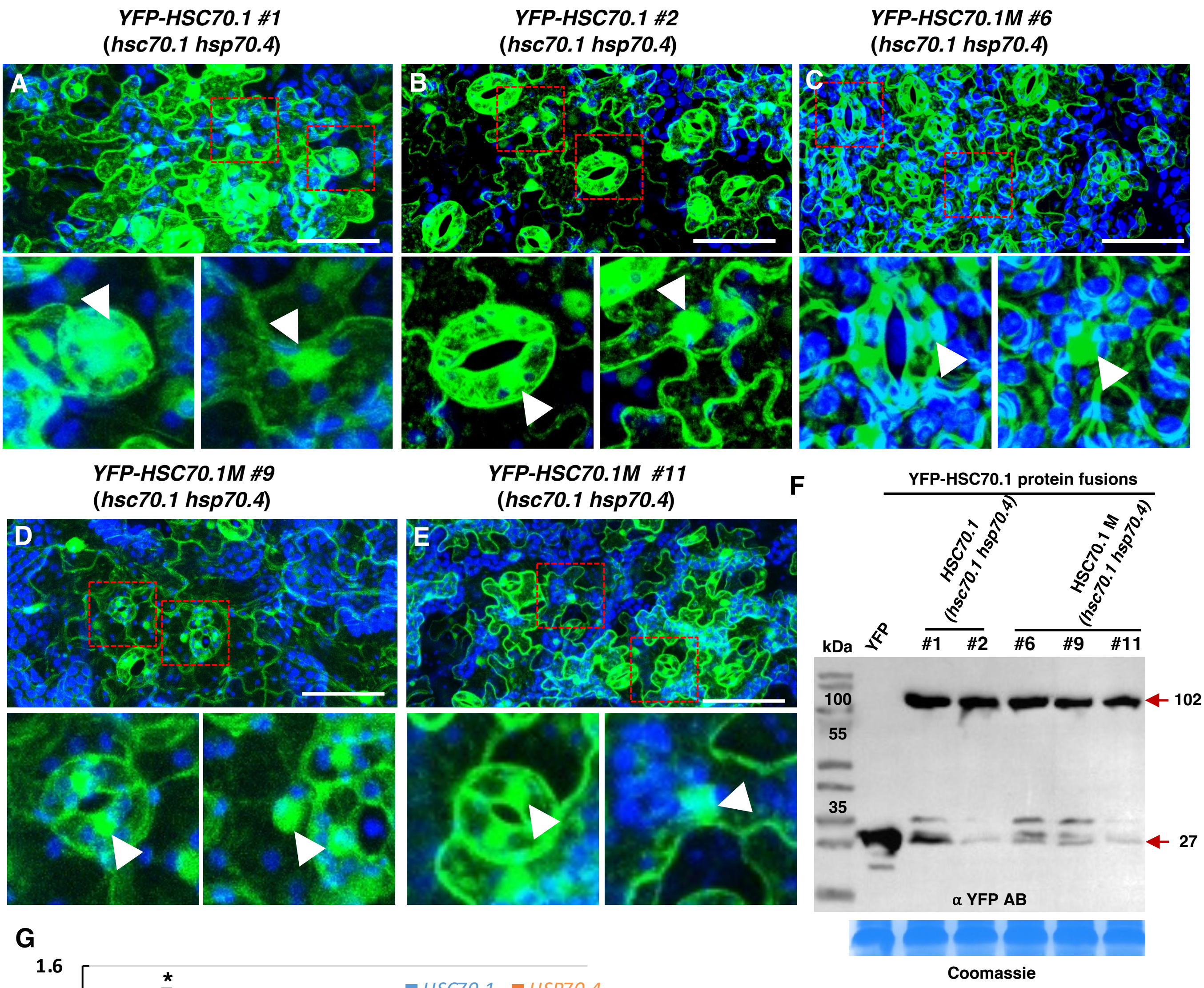


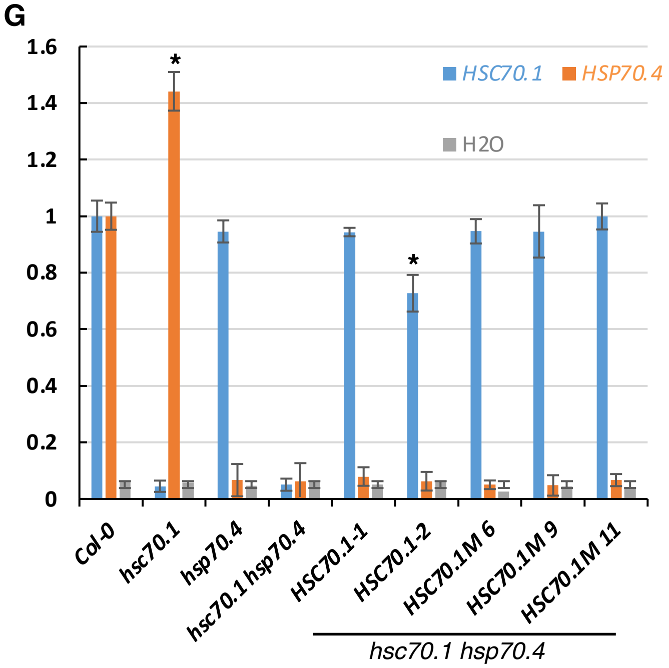


**Figure S9. Expression of fusion constructs in *hsc70.1 hsc70.4* mutants.**

A to E, Confocal laser scanning microscopy (CLSM) images of transgenic *hsc70.1 hsp70.4* lines. All YFP fusion proteins were detected in both cytoplasm and nuclei. Blue color: plastid auto-fluorescence. Green color: YFP fluorescence. Bar: 10 $\mu$m.

F, Western-blot detection of YFP-HSC70.1 and YFP-HSC70.1M fusion proteins produced in two-week-old transgenic *hsc70.1 hsp70.4* mutant. Lower panel: loading control. Red arrows indicated expected sizes.

G, qRT-PCR assays on independent two *YFP-HSC70.1* and three *YFP-HSC70.1M* transgenic *hsc70.1 hsp70.4* lines used for phenotyping. Except line #2 all show no significant differences in their *HSC70.1* transcript levels. Bars: SE.

### Supplementary Figure S10
### Yang et al. 2022

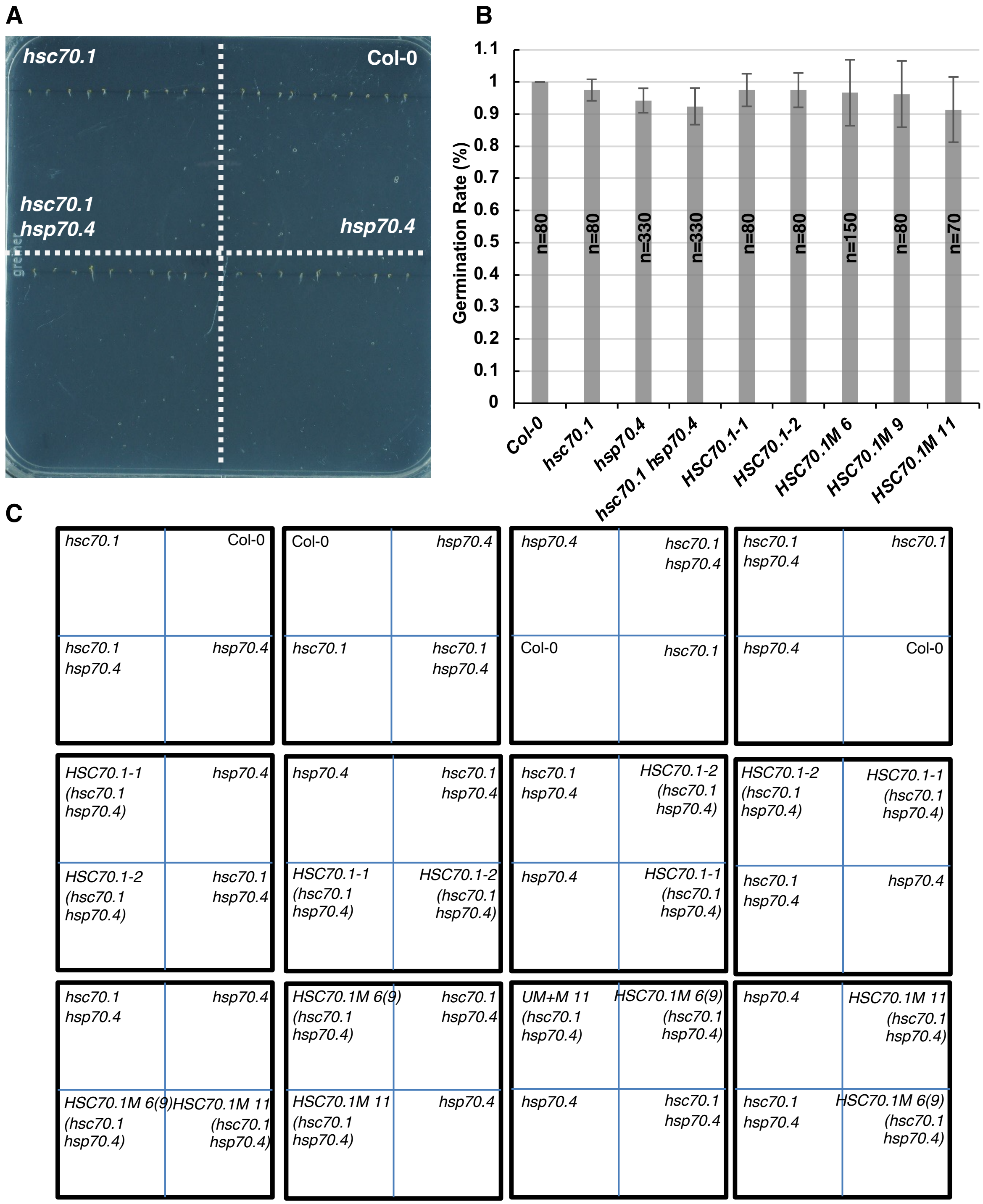


**Figure S10. Germinating seeds analysed for root growth.**

A, Representative image of used seeds vertically germinate on agar medium at two days. A seed was considered germinating when radicle protrusion was visible.

B, Germination of wild-type (Col-0), *hsc70.1, hsp70.4, hsc70.1 hsp70.4, HSC70.1-1 (hsc70.1 hsp70.4), HSC70.1-2 (hsc70.1 hsp70.4), HSC70.1M-6 (hsc70.1 hsp70.4), HSC70.1M-9 (hsc70.1 hsp70.4),* and *HSC70.1M-11 (hsc70.1 hsp70.4)* seeds. T-test analysis indicates no significant germination difference among the seeds. The seeds were stratified at 4°C in the dark for 2 days and then grow them on the plates under long day condition. n = number of seeds analyzed; Error bar: ± SE; Significance was calculated using Student's T-test (two tails).

C, The distribution scheme of different genotypic seeds on the plates to avoid position biased growth.

### Supplementary Figure S11
### Yang et al. 2022

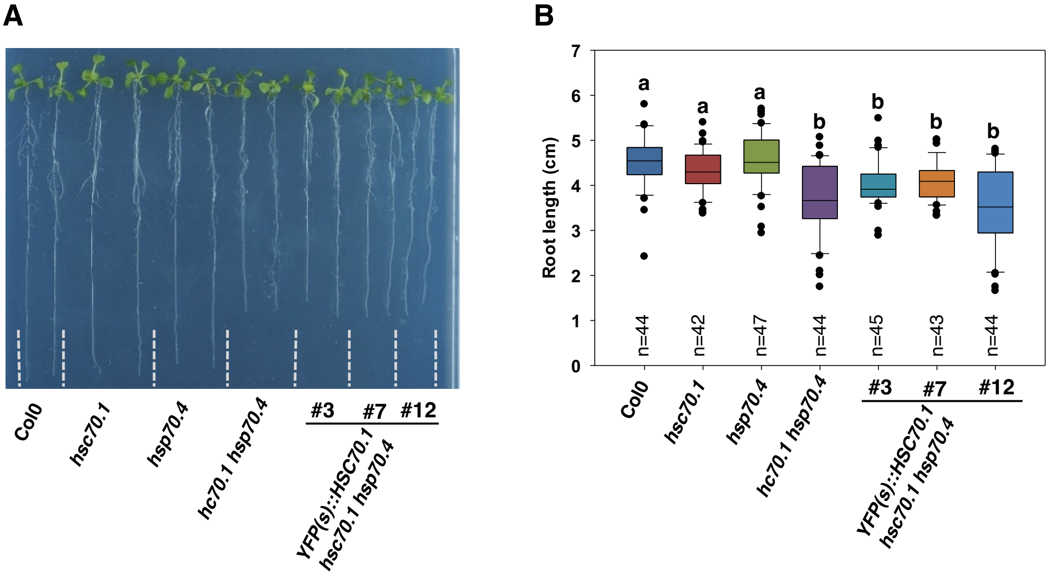


**Figure S11. *YFP(s)::HSC70.1* transcript and *hsc70.1 hsp70.4* root growth.**

A, Representative pictures of analyzed wild-type (Col-0), *hsc70.1*, *hsp70.4*, *hsc70.1 hsp70.4*, *YFP(s)::HSC70.1 #3* (*hsc70.1 hsp70.4), YFP(s)::HSC70.1 #7* (*hsc70.1 hsp70.4*) and *YFP(s)::HSC70.1 #12* (*hsc70.1 hsp70.4)* plants 14 days after germination.

B, Quantitative data of measured primary root length of wild-type and indicated mutant plants. Box plot graph: Boxes indicate variation between datasets and means; n = number of analyzed plants; error bar: ± SE; black dots: measurements out of ± SE range. Significance was calculated using Student's T-test (two tails); p value indicated by a and b: a, b<0.001.

### Supplementary Figure S12
### Yang et al. 2022

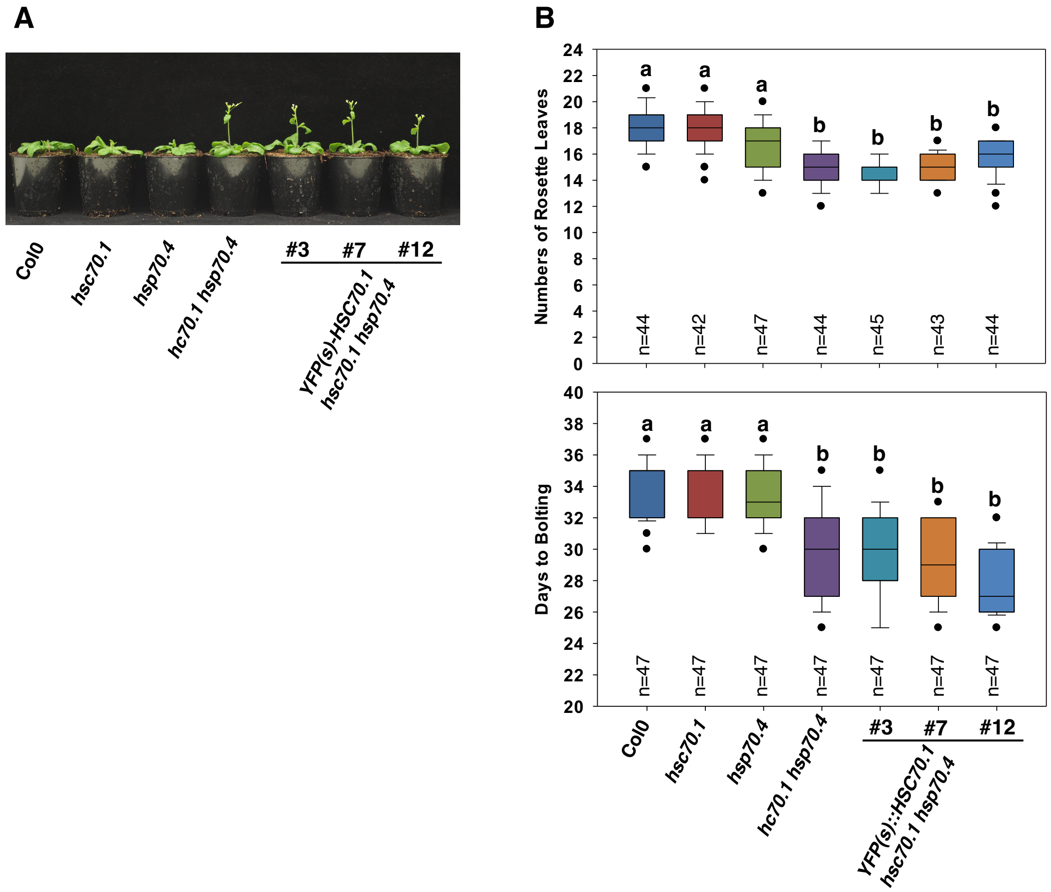


**Figure S12. *YFP(s)::HSC70.1* fusion transcript and lack of *hsc70.1 hsp70.4* early flowering phenotype complementation**.

A, Representative pictures of analyzed wild-type (Col-0), *hsc70.1*, *hsp70.4*, *hsc70.1 hsp70.4*, *YFP(s)::HSC70.1 #3* (*hsc70.1 hsp70.4), YFP(s)::HSC70.1 #7* (*hsc70.1 hsp70.4*) and *YFP(s)::HSC70.1 #12* (*hsc70.1 hsp70.4)* plants 30 days after germination.

B, Upper panel: age of wild-type and mutant plants at bolding. Lower panel: Numbers of rosette leaves at time of bolding of wild-type and indicated mutant plants. Box plot graph: Boxes indicate variation between datasets and means; n = number of plants analyzed; error bar: ± SE; black dots: measurements out of range ± SE; significance was calculated using Student's T-test (two tails); p value indicated by a and b: a, b<0.001.

### Supplementary Table S1
### Yang et al. 2022

**Table S1. Western blot bands density relative to Mock control in all three repeated independent experiments.**

| Minutes | YFP | | | YFP-HSC70.1 | | | YFP-HSC70.1 ΔSVR | | |
|---|---|---|---|---|---|---|---|---|---|
| | Replicate1 | Replicate2 | Replicate3 | Replicate1 | Replicate2 | Replicate3 | Replicate1 | Replicate2 | Replicate3 |
| 0 | 1.00 | 1.00 | 1.00 | 1.00 | 1.00 | 1.00 | 1.00 | 1.00 | 1.00 |
| 15 | 1.08 | 1.01 | 1.01 | 1.53 | 1.56 | 1.43 | 1.02 | 1.02 | 1.05 |
| 30 | 1.09 | 1.06 | 1.09 | 1.58 | 1.68 | 1.59 | 0.93 | 1.02 | 1.00 |
| 60 | 1.10 | 1.05 | 0.99 | 1.01 | 1.07 | 0.98 | 0.90 | 1.05 | 0.99 |

### Supplementary Table S2
### Yang et al. 2022

**Table S2. *A. thaliana HSC70* orthologous transcripts annotated as mobile.**

| Name | TAIR gene number | Mobility (Thieme et al. 2015) | Annotation |
|---|---|---|---|
| **HSC70.1** | AT5G02500 | A.th/Cuscuta | ARABIDOPSIS THALIANA HEAT SHOCK COGNATE PROTEIN 70-1, AT-HSC70-1, ATHSP70-1, HEAT SHOCK COGNATE PROTEIN 70, HEAT SHOCK COGNATE PROTEIN 70-1, HEAT SHOCK PROTEIN 70-1, HSC70, HSC70-1, HSP70-1 |
| HSC70.2 | AT5G02490 | n/a | ATHSP70-2, HSP70-2 |
| **HSC70.3** | AT3G09440 | A.th/Cuscuta | Heat shock protein 70 (Hsp 70) family protein |
| HSC70.4 | AT3G12580 | n/a | ARABIDOPSIS HEAT SHOCK PROTEIN 70, ATHSP70, HEAT SHOCK PROTEIN 70, HSC70-4, HSP70 |
| **HSC70.5** | AT5G09590 | A.th/Cuscuta | HEAT SHOCK COGNATE, HSC70-5, MITOCHONDRIAL HSO70 2, MTHSC70-2 |
| **HSC70.6** | AT4G24280 | A.th/Cuscuta&PED/COL | CHLOROPLAST HEAT SHOCK PROTEIN 70-1, CPHSC70-1 |
| **HSC70.7** | AT5G49910 | A.th/Cuscuta&PED/COL | CHLOROPLAST HEAT SHOCK PROTEIN 70-2, CPHSC70-2, HEAT SHOCK PROTEIN 70-7, HSC70-7 |
| HSC70.8 | AT2G32120 | n/a | HEAT-SHOCK PROTEIN 70T-2, HSP70T-2 |
| **HSC70.9** | AT4G37910 | A.th/Cuscuta | MITOCHONDRIAL HEAT SHOCK PROTEIN 70-1, MTHSC70-1 |
| HSC70.10 | AT1G16030 | n/a | HEAT SHOCK PROTEIN 70B, HSP70B |
| HSC70.11 | AT5G13820 | n/a | ATBP-1, ATBP1, ATTBP1, H-PROTEIN PROMOTE, HPPBF-1, TBP1, TELOMERIC DNA BINDING PROTEIN 1 |
| HSC70.12 | AT4G32208 | n/a | heat shock protein 70 (Hsp 70) family protein |
| HSC70.13 | AT1G09080 | n/a | BINDING PROTEIN 3, BIP3, Heat shock protein 70 (Hsp 70) family protein |
| **HSC70.14** | AT1G79930 | A.th/Cuscuta | ATHSP70-14, HEAT SHOCK PROTEIN 91, HSP91 |
| **HSC70.15** | AT1G79920 | A.th/Cuscuta | ATHSP70-15, HEAT SHOCK PROTEIN 70-15, HSP70-1 |
| **HSC70.16** | AT1G11660 | PED/COL | HEAT SHOCK PROTEIN 70-16, HSP70-16 |
| **HSC70.17** | AT4G16660 | A.th/Cuscuta&PED/COL | HEAT SHOCK PROTEIN 70, HSP7 |
| HSC70.18 | AT1G56410 | n/a | EARLY-RESPONSIVE TO DEHYDRATION 2, ERD2, HEAT SHOCK PROTEIN 70T-1, HSP70T-1 |

*A.th* / Cuscuta indicates mobile from *A.th* to cuscuta; *PED* / Col-0 indicates mobile in grafted Arabidopsis plants; n/a: not analysed.

### Supplementary **Table S3**
### **Yang et al. 2022**

**Table S3. The chemical reactions, the system of ordinary differential equations and all parameters in the model.**

The default parameters for the model are given below. For the system without feedback, the kinetic on-rate of HSC70.1 on its mRNA was set to $k_{+h.H} = 0$, and the translation rate, $\alpha_{H,1}$, was adjusted to $\alpha_{H,1} = 0.30394797$, to give the same steady state protein level of HSC70.1 as the system with feedback. The initial concentration of HSC70.1 was set to zero, except for simulating the in vitro translation experiments for which a range of different started values were used to mimic the experiment. $\alpha_M = 300$ is only initiated after a defined time (t=200 in the simulations for Figure 3E).

| Process | Chemical equation | Symbols | Comments |
|---|---|---|---|
| **HSC70.1 translation from *HSC70.1* mRNA** | h + R ⇔ h + R + H | h = HSC70.1 mRNA<br>R = ribosomes<br>H = HSC70.1 protein | The rate depends on the availability of H;<br>Total amount of h, $h_T$, constant (see below) |
| **Negative autoregulation of HSC70.1 translation** | h + H ⇔ h.H | h.H = HSC70.1 mRNA-protein complex | h.H cannot be translated |
| **HSC70.1 guiding refolding of mis-folded proteins** | H + M ⇔H.M -> H + C | M = mis-folded protein<br>C = correctly folded protein<br>H.M = HSC70.1 protein complex with misfolded protein | Mis-folded proteins only produced under stress |
| **Protein degradation** | H -> 0<br>M -> 0<br>C -> 0 | | Amount of ribosomes assumed constant and not modelled explicitly. |

| **Ordinary differential equations** | Parameters (a.u.) |
|---|---|
| $\frac{dH}{dt} = \alpha_H(h_T - h) - k_{+h.H}H(h_T - h) + k_{-h.H}h.H - k_{H.M}HM + k_{H,C}H.M - \delta H$<br>$\frac{dh.H}{dt} = k_{+h.H}H(h_T - h) - k_{-h.H}h.H$<br>$\frac{dM}{dt} = \alpha_M - k_{H.M}HM - \delta M$<br>$\frac{dH.M}{dt} = k_{H.M}HM - k_{H,C}H.M$ | $h_T = 100$,<br>$\alpha_H = \alpha_{H,0} + \alpha_{H,1}H/(H+K)$, K = 100,<br>$\alpha_{H,0} = 0.01$, $\alpha_{H,1} = 1$<br>$k_{+h.H} = 1$, $k_{-h.H} = 100$,<br>$\delta = 0.1$, $k_{H.M} = 0.01$,<br>$k_{H.C} = 0.1$, $\alpha_M = 300$ |
